# InterOpt: Improved gene expression quantification in qPCR experiments using weighted aggregation of reference genes

**DOI:** 10.1101/2023.04.24.538126

**Authors:** Adel Salimi, Saeid Rahmani, Ali Sharifi-Zarchi

## Abstract

Quantification of gene expression is a crucial task in biomedical studies. Although high-throughput methods enable rapid and simultaneous expression quantification of coding or non-coding regions, qPCR is still vastly used due to its high availability, sensitivity, specificity, reproducibility, low cost, and ease of use. A limitation of qPCR has been the need of internal controls or reference genes (RGs) with stable expression in different conditions, to normalize the expression level of the other target genes. So far, several stability criteria and numerous methods for selecting a group of RGs have been proposed, however, important challenges must be addressed. Here we introduce a mathematical basis for precise modeling of qPCR expression normalization and justify widely used stability measures of RGs. We then propose a family of scale-invariant functions, as an alternative to the geometric mean, to optimize aggregated expression of RGs. We provide closed-form optimizations for several scale-invariant aggregation functions. Among them, we show the superiority of weighted geometric mean, whose parameters optimize standard deviation as the stability measure of aggregated RGs expression. We provide experimental support for this finding using real data of solid tumors and liquid biopsies of different sample sizes.

The proposed methods can be easily integrated in the existing qPCR expression normalization pipelines of genes and non-coding RNAs. We also provide an implementation of the proposed methods as an R package, with GPU acceleration.

**Availability and implementation:** https://github.com/asalimih/InterOpt

## 1 INTRODUCTION

Reverse transcription quantitative PCR (RT-qPCR) is one of the most utilized techniques to quantify RNA molecules. Despite high-throughput methods such as RNA-seq are widely used for expression quantification, qPCR is still the primary molecular diagnostic test for clinical and research purposes due to its high specificity and sensitivity, low cost, and reproducibility (Saiki *et al*., 1988; Garibyan and Avashia, 2013). There are, however, different sources of technical errors that make the accuracy and power of qPCR highly dependent on the normalization procedure.

The qPCR method uses repeated cycles of DNA amplification to measure the expression of target gene(s) in a given sample. The amount of the target region approximately doubles during each amplification cycle. We define *Cycle Threshold* (CT) as the first cycle the amount of amplified target region exceeds a fixed threshold (Acinas *et al*., 2005). This raw CT value is affected by two sources of variation: biological and technical (Taylor *et al*., 2019). To accurately measure expression alternation of a target gene among different conditions, we need to minimize these variations. For this purpose, cross-sample normalization is performed to make the expression levels of a target gene comparable among different samples, which leads to statistically authentic results. The most usual way of cross-sample normalization employs a *reference gene* (RG) and normalizes the expression of each target gene by subtracting the expression of RG from it. The assumption behind these techniques is that the RG expression is unaffected in different conditions (Vandesompele *et al*., 2002). Choosing an unstable or differentially expressed RG could lead to contradictory results. For example, if an RG has expression alternation between the treatment vs. control groups, a correlated target gene would show lower or no significant expression change after normalization with this RG. An example is provided in (Ghanbari *et al*., 2021).

An optimal RG is a gene with minimal biological variation among different samples, which is also highly expressed in all samples. There are housekeeping genes (e.g. GAPDH) that are widely used for this purpose. Several studies, however, have shown those so-called housekeeping genes have high variation of expression in specific tissues or diseases (Altenberg *et al*., 2004; Guo *et al*., 2013). Moreover, some studies question the existence of any housekeeping gene with above-mentioned conditions (Zhang *et al*., 2015). This challenge is even more apparent in non-coding RNA studies, particularly when using circulating miRNAs as cancer biomarkers (Marabita *et al*., 2016; Rice *et al*., 2015). Therefore, using multiple and study-specific RGs is highly recommended in the standard guidelines (Bustin *et al*., 2009; Taylor *et al*., 2019).

Using multiple RGs in a qPCR experiment raises several challenges, including the need to aggregate expression levels of multiple RGs into a single reference value for normalizing the other genes. This single value can be considered as the expression level of a virtual RG. Additionally, we need to measure the expression *stability* of a real or virtual RG which is maximized when that RG shows zero expression variation among different conditions. Although the later challenge is well studied (Vandesompele *et al*., 2002; Andersen *et al*., 2004; Sundaram *et al*., 2019), there is a paucity of literature about the former one which is the main focus of this paper.

The standard deviation (SD) of CT values is one of the main stability measures of an RG. In the case of 2 candidate RGs with equal SD, the one with a higher expression level is preferable. The reason is genes with low abundance have a higher chance of not being detected due to technical errors. Accordingly, the coefficient of variation (CV) is another widely used measure for an RG’s stability, which is the SD of expression divided by mean expression. Both SD and CV are vastly used to measure the stability of an RG in the literature, but no theoretical explanation has been provided (Li *et al*., 2019; Sundaram *et al*., 2019; Li *et al*., 2020).

In this work, we first elucidate the theoretical assumptions behind SD and CV as RG expression stability measures. Next, by introducing the family of scale-invariant functions, we explain the reason arithmetic and geometric mean functions can be used for the aggregation of multiple RGs. Then by providing mathematical solutions for the optimization of the weighted version of geometric and arithmetic mean functions, we present four novel weighted aggregation methods to optimally minimize the SD and CV of a virtual RG. Each proposed method is defined by an aggregation function (geometric or arithmetic mean) and a measure (SD or CV) to be minimized, and we call them weighting methods. To evaluate these methods, we utilized qPCR array datasets. qPCR array datasets with a high number of genes or other RNA molecules can be normalized without RGs. This feature enabled us to design a benchmarking pipeline which we could calculate stability measures of different combinations of weighted RGs on the normalized data and compare the weighting methods. Finally, we chose the best method (geom(sd)) and showed the weights of the weighted geometric mean optimized on SD of logarithm of expression can be calculated solely from the raw CT values without being affected by the gene-independent technical variations. Experimental evaluations show that this method can also be used in low sample size conditions.

## 2 RELATED WORKS

Various RG expression stability methods have been previously proposed and they’re being highly utilized for the selection process of RGs and their stability assessment.

GeNorm is an iterative method that uses pairwise variation to measure the stability of RGs. In each iteration the candidate with the worst stability score is removed. Then, the procedure is repeated until only 2 genes remain. the method’s assumption is all input genes should have low expression variation. (Vandesompele *et al*., 2002). NormFinder is another widely used tool that suggests a mathematical model that separates technical and biological variations, then eliminates the technical biases to find the RG with the lowest biological variation (Andersen *et al*., 2004). NormiRazor offers a GPU-based implementation of GeNorm, NormFinder and BestKeeper to examine the stability of high number of RG combinations in parallel (Grabia *et al*., 2020).

To our knowledge there has previously been only one study that has approached the problem of aggregating multiple reference genes. This method uses weighted geometric mean and follows a heuristic approach to define each RG’s weight by the ratio of their standard deviation (Qureshi and Sacan, 2013). We have named it geom(sd r), and it is been evaluated along with our proposed weighting methods.

## 3 MATERIALS AND METHODS

This section is composed of six parts. In 3.1, we review SD and CV as stability criteria for an optimal RG by modeling the normalization procedure and gene expression distribution. Next, in 3.2, we show why arithmetic and geometric mean could be used as aggregation functions to build an virtual RG from multiple RGs. In 3.3, we propose solutions to optimize the weighted version of those functions based on different stability criteria. Additionally detailed description of all the proposed methods can be found in the appendix. In 3.4, we present a benchmarking pipeline to evaluate the proposed weighting methods in different biological situations as well as number of samples and the obtained datasets are explained in 3.5. Finally the implementation and availability details of proposed methods and statistical analyses are described in 3.6. It is worth mentioning that in this study, a gene consists of both coding and none-coding genes, and by gene expression in a qPCR study, we mean RNA concentration which is affected by both gene transcription and degradation.

### 3.1 The criteria of optimal reference gene

A widely used application of gene expression quantification is differential expression analysis. In which genes expression are compared among samples using fold change or ratio:

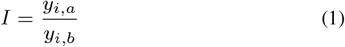

Here *y*_*i,a*_ is the expression of gene *i* in sample *a* and *y*_*i,b*_ is the expression of gene *i* in sample *b*. We aim to find the biological variations however, the CT values of a qPCR experiment also comprise technical variations. One of the primary sources of technical variation is the different amounts of initial RNA concentration at the start of the qPCR process for each sample. We can model this effect as a coefficient for each sample’s different genes:

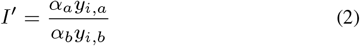

Here *α*_*a*_*y*_*i,a*_ and *α*_*b*_*y*_*i,b*_ are the raw measured concentration of gene *i* in samples *a* and *b* accordingly and *α*_*a*_ and *α*_*b*_ represent the technical variation as coefficients. In order to remove this technical variation, each gene expression is divided by an RG (*y*_*r*_) which is also affected by the technical variation:

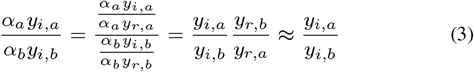

To find the true ratio of the target gene, the ratio of the RG expression in different samples should be close to 1. *Z* is a continuous random variable with probability density function *f*_*Z*_ (*z*; *θ*) representing this ratio. Hence the objective can be defined as maximizing the probability density of *Z* in the proximity of 1:

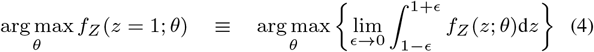

*θ* represents the parameters of the probability density function.

#### 3.1.1 Gaussian expression distribution leads to CV as RG stability measure

One of the common distributions to model gene expression is the Gaussian distribution. If we model the expression of an RG *r* by a Gaussian distribution with mean *μ* and standard deviation *σ*, the distribution of the ratio of the gene in two different samples would be Eq.6 (Díaz-Francés and Rubio, 2013).

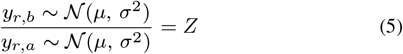

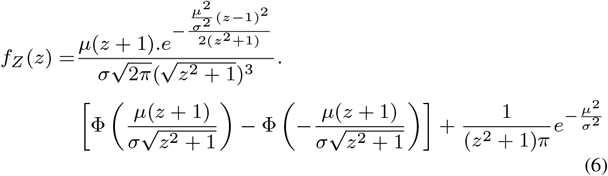

The definition of coefficient of variation of a gene is defined as standard deviation of gene expression divided by the mean expression:

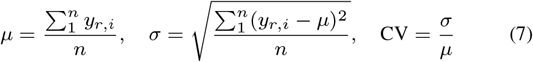

Where *n* is the number of samples. Considering the Eq.4, it can be proven that maximization of 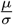 is equivalent to maximizing *f*_*Z*_ (*z* = 1; *μ, σ*) or in other words minimizing the coefficient of variation (CV) of the Gaussian distribution. This is detailed in Theorem 2 of the appendix. This implies CV can be used as a measure of stability, given that the distribution of the RG expression follows a Gaussian distribution.

#### 3.1.2 Log-normal expression distribution leads to SD as RG stability measure

Another previously suggested distribution to model gene expression is log-normal distribution (Bengtsson *et al*., 2005). As the expression of a gene is always a positive number, this distribution has some benefits compared to the Gaussian distribution.

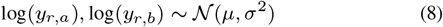

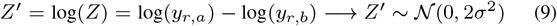

Now we can rewrite Eq.4 in terms of *Z*′:

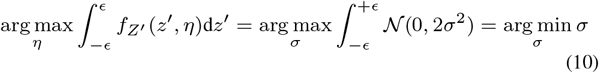

In conclusion, assuming gene expression follows a log-normal distribution, an RG with lower SD of the logarithm of expression is more stable.

### 3.2 Geometric and Arithmetic Mean as Aggregation Functions

We model gene independent technical variations in the form of scaling operations in each sample. As explained in Eq.3, an RG should preserve these variations. Therefore the aggregated virtual RG of a set of *d* RGs should also follow the same rule:

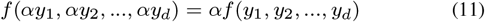

We call the Eq.11 family of functions, scale-invariant functions. Arithmetic and geometric mean are members of this family of functions. Their weighted counterparts are also scale-invariant functions and their weights can be optimized depending on the objective.

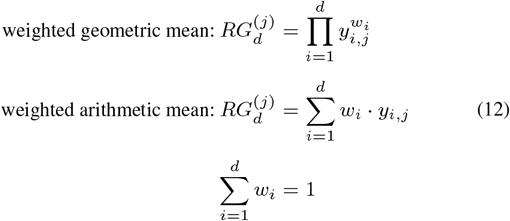

Here 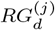 is a weighted mean of a combination of *d* RGs in the sample

*j. y*_*i,j*_ is the expression value of *i*-th RG (among the *d* RGs) in the sample *j* and *w*_*i*_ is the *i*-th RG weight. Throughout the paper, *RG*_*d*_ is also referred as a virtual RG resulted from aggregation of d RGs.

### 3.3 Optimizing Weighted Geometric/Arithmetic Mean

Weighted geometric and arithmetic mean aggregation functions are parametric and the weights can be optimized based on an objective function. Taking together all combinations of arithmetic and geometric mean as the aggregation functions and SD and CV as stability measures we introduce the following weighting methods:

- geom(sd): Aggregation of *d* RGs using weighted geometric mean, optimized on SD of logarithm of *RG*_*d*_.
- geom(sd+): Improved version of geom(sd) for small sample size datasets
- arith(cv): Aggregation of *d* RGs using weighted arithmetic mean, optimized on CV of *RG*_*d*_.
- geom(cv): Aggregation of *d* RGs using weighted geometric mean, optimized on CV of *RG*_*d*_.
- arith(sd): Aggregation of *d* RGs using weighted arithmetic mean, optimized on SD of logarithm of *RG*_*d*_.

Each of these weighting methods and their optimization solutions are described in the next sections.

#### 3.3.1 geom(sd)

This section determines the optimal weighted geometric mean to minimize the SD of the logarithm of the aggregated RG. The geometric mean is equivalent to the arithmetic mean in the logarithmic space; therefore the optimization problem would be as follows:

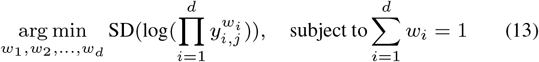

In Eq.13, *d* is the number of RGs, and *y*_*i,j*_ is the expression of the *i*-th RG in sample *j*. by applying the logarithm to the product and injecting the constraint into the equation, we can simplify it as Eq.14 (details in Theorem 4 of the Appendix):

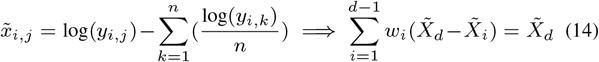

Here *n* is the sample size and 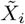 is the mean centered expression of the *i*-th RG in logarithmic scale. The closed form solution of Eq.14 is as follows:

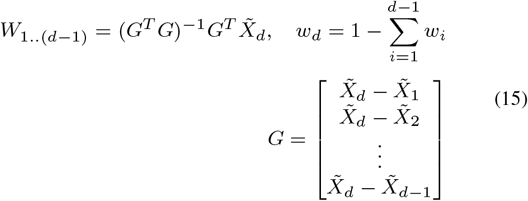

*W*_1..(*d*−1)_ is a 1 *×* (*d* − 1) matrix consisting of the first (*d* − 1) elements *W*

#### 3.3.2 geom(sd+)

Eq.13 can also be formulated using random variables. Suppose *X*_1_ and *X*_2_ are two random variables representing the logarithm of two RGs expression. The variance of their weighted average would be:

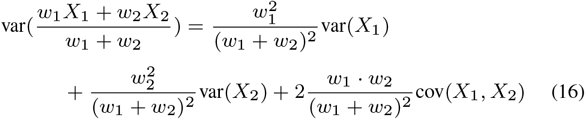

Setting the derivative of Eq.16 to zero with respect to each of the weights leads to a simplified solution:

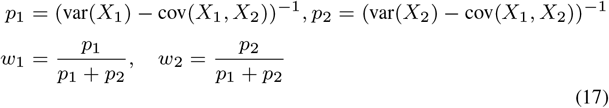

Eq.17 only requires to estimate the variance and covariance of the RGs expression. This enabled us to enhance geom(sd) by utilizing specialized covariance matrix estimation methods for small sample size datasets. After comparing covariance estimation methods for different sample sizes (n), we employed a hybrid approach that uses the oracle approximation shrinkage method for *n <* 15 (Chen *et al*., 2010), soft thresholding for 15 ≤ *n <* 85, and hard thresholding for *n* ≥ 85 (Bickel and Levina, 2008). This hybrid method is named geom(sd+) throughout this paper.

The CT values obtained from qPCR are subject to technical and biological variations. Assuming the technical variation be gene-independent, it can be proven that by modeling the technical variation as an additive random variable *F* and we replace *X*_1_ and *X*_2_ with *X*_1_ + *F* and *X*_2_ + *F*, the solution would still be Eq.17 (details are provided in theorem 7 of appendix). This shows the robustness of geom(sd) and geom(sd+) method to gene-independent technical variations of qPCR CT values.

#### 3.3.3 arith(cv)

*Y* is a *d × n* matrix containing the expression of *d* RGs in *n* samples. Here the goal is to minimize the CV of the weighted arithmetic mean of the RGs. This optimization problem is demonstrated in Eq.18:

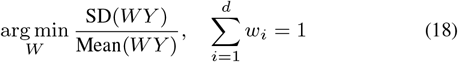

This equation can be written as:

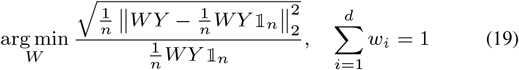

where 𝟙_*n*_ is a *n* × 1 matrix of ones.

Surprisingly a closed form solution for this problem can be obtained (details are provided in theorem 5 of appendix):

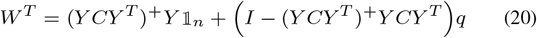

Where *C* is the Centering Matrix 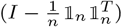, *H*^+^ denotes the pseudo-inverse of *H, I* is the identity matrix, and *q* is an arbitrary vector.

#### 3.3.4 geom(cv)

The matrix product form of weighted geometric mean function is exp(*W* ln(*Y*)) So if *Y* ′ = ln(*Y*), then the optimization problem would be:

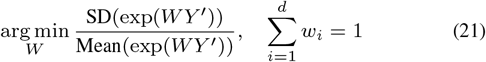

Eq.21 can be rewritten as:

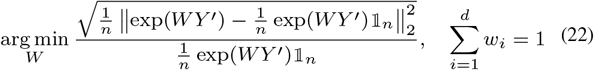

1_*n*_ is a *n* 1 matrix of ones.

Contrary to the previous method, there is no closed form solution here. However, the gradient with respect to *x* where 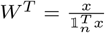 can be obtained.

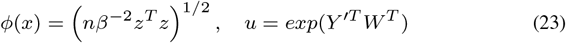

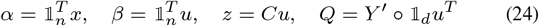

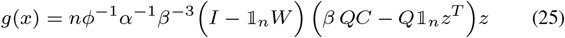

Where *g*(*x*) is the gradient of the loss function with respect to *x*. We used a stabilized version of the Barzilai-Borwein method (Fletcher, 2005; Burdakov *et al*., 2019) to optimize it (details are provided in theorem 6 of appendix).

#### 3.3.5 arith(sd)

Unlike previous methods no mathematical solution was provided for this optimization problem. Thus, based on the constraint that weights sum to 1, a numerical procedure in which an exhaustive search through weights with a precision of 0.01 was used.

### 3.4 Benchmark

In order to evaluate and compare the weighting methods on real data, we acquired qPCR array datasets and designed the benchmark workflow depicted in Fig1. First lowly expressed and genes with none-detects in more than 4 samples are removed. The remaining none-detects were imputed using the *nondetects* R package (McCall *et al*., 2014). Next, depending on the weighting method, for any combination of *d* genes, the Eq.12 is calculated, and the resulting virtual RG’s stability is measured based on SD, CV, GeNorm and NormFinder. The individual stability measures are then converted to their corresponding standard z-scores, and their average is taken to obtain the aggregated Stability measure.

**Fig. 1:**
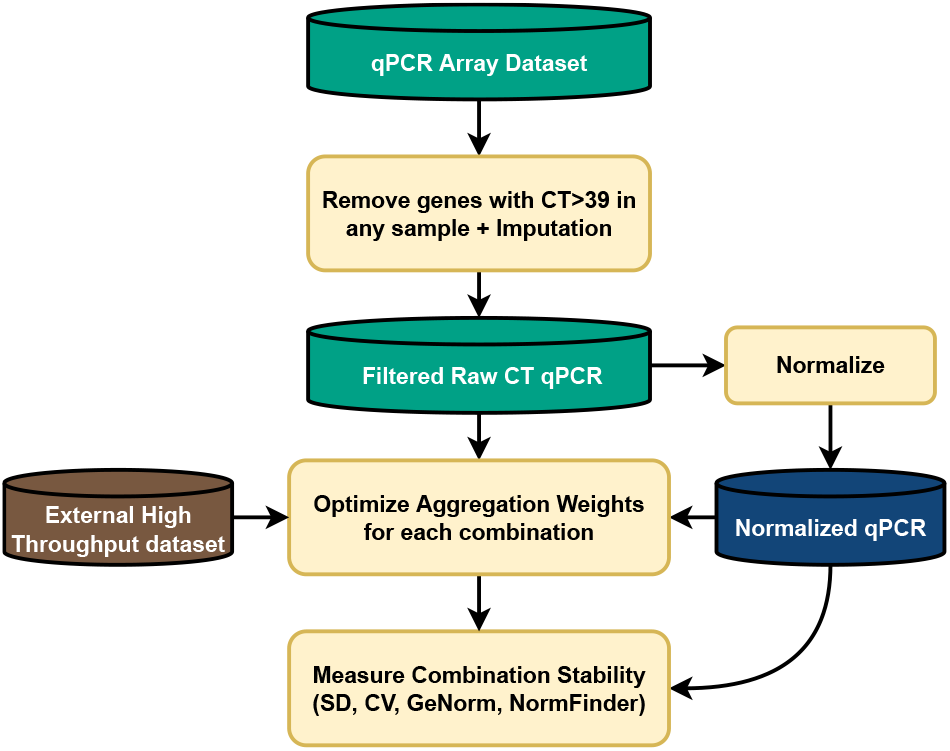
Benchmark workflow. SD: Standard Deviation, CV: Coefficient of Variation

The weight optimization of the weighting methods is executed on three different types of data. Raw CT values, normalized CT and a normalized external high-throughput dataset.

### 3.5 High-Throughput and qPCR Datasets

Two qPCR array datasets (GSE78870 and GSE50013) were obtained from the Gene Expression Omnibus (GEO). The GSE78870 contained the expression of 768 miRNAs in 106 primary breast cancer specimens, and GSE50013 contains the expression of 762 (258 detectable) miRNAs in the plasma of 20 patients with hepatocellular carcinoma, as well as 20 healthy donors. To normalize the qPCR array datasets, we have adopted global normalization (Mestdagh *et al*., 2009). In this method the mean expression value of all expressed microRNAs (CT*>*35) in each given sample is used as a normalization factor.

The Cancer Genome Atlas (TCGA) is a comprehensive cancer genomics program that includes molecular datasets for a different types of cancer tissues (Weinstein *et al*., 2013). Using the TCGAbiolinks package, the breast cancer miRNA-Seq profile of 1097 tumor samples were obtained from the TCGA portal. The count matrix was normalized in count per million (CPM).

For the experimental validation, we used the qPCR dataset from our previous study containing 12 pairs of breast cancer and adjacent normal tissue (Ghanbari *et al*., 2021).

### 3.6 Implementation and Statistical Analysis

All the proposed weighting methods were implemented in an easy to use open source R package called InterOpt which is available at (https://github.com/asalimih/InterOpt). geom(sd) and arith(cv) were implemented based on their closed form solutions. For geom(cv) a stabilized version of the Barzilai-Borwein gradient method was utilized. It requires less computation and greatly speeds up the convergence compared to other gradient methods. Since no solution for the arith(sd) method was given, an exhaustive search through weights was utilized as an alternative approach.

Running the proposed benchmark for thousands of combinations is not trivial on a single CPU core. Thus we utilized a CUDA accelerated implementation of stability measures (GeNorm, NormFinder) called NormiRazor (Grabia *et al*., 2020). We modified it to integrate aggregation weights and also be capable of handling combinations of *d >* 3 number of RGs.

All statistical analyses were executed in the RStudio integrated development environment and R language v3.6.1. Paired Wilcoxon rank sum test was carried out for stability comparison between weighting methods with a p.value significance level of 0.01. Covariance estimation of geom(sd+) method was carried out by the CovTools v0.5.4 package (https://github.com/kisungyou/CovTools) (Lee and You, 2021). Figures were produced using ggplot2 v3.3.5 (Wickham, 2016), pheatmap v1.0.12 (Kolde, 2019), cowplot v1.0.0 (Wilke, 2020), ggsci v2.9 (Xiao, 2018) and ggsignif v0.6.3 Constantin and Patil (2021) packages. An open source GitHub repository for reproducing all the results and figures is also available at (https://github.com/asalimih/InterOpt-paper)

## 4 RESULTS

Following the benchmark presented in Fig.1, our validation for the proposed methods was carried out in three scenarios which are different based on the data type that the aggregation weights were calculated from: It is important to note that the data on which the weights are optimized

- Raw CT values of the qPCR array: shows the effect of qPCR technical variations on the performance of weighting methods.
- Normalized qPCR array: exhibits how much the weighting method could lower the biological variation if the data had very small to no technical variation. The mean CT of all expressed microRNAs (CT*<*35) is used as the normalization factors.
- An external biologically compatible RNA-Seq dataset: shows if the weights could be calculated from a separate high-throughput dataset.

In all three scenarios the stability measures (SD, CV, GeNorm and NormFinder) were calculated on the normalized qPCR array but the weights of the weighting methods were optimized on the aforementioned data types.

### 4.1 Stability Comparison of Weighting Methods

Here we utilized two qPCR array datasets with different variability levels across samples. A breast cancer tissue dataset and a liver cancer plasma dataset with median expression SD of 1.6 and 2.77 respectively among their miRNAs. The standard deviation of the log2 of the normalization factors was considered as an estimate of the technical variation caused by RNA abundance of the samples which were 0.45 and 0.72 for the breast and liver cancer datasets, respectively. As illustrated in Fig.2A and Fig.2B, on both datasets the geom(sd) method outperforms all other methods in terms of the aggregated stability, specifically when the aggregation weights are calculated based on the raw CT values. The geom(cv) method is the second-best method on the breast cancer dataset, but it has the worst stability on the liver cancer dataset. By contrast, the arith(sd) method performs as well as geom(sd) in the liver dataset. The difference between the normalized and raw CT results suggests that geom(cv) and arith(cv) are sensitive to technical variations in raw CT values, and thus not suitable for datasets with high technical variation. All methods tested outperform the usual geometric mean. Table1 and Table2 show each stability measure separately. Expectedly, geom(cv) had a lower CV than others, but high numbers on other measures have made its overall stability worse than others in the liver dataset.

**Fig. 2:**
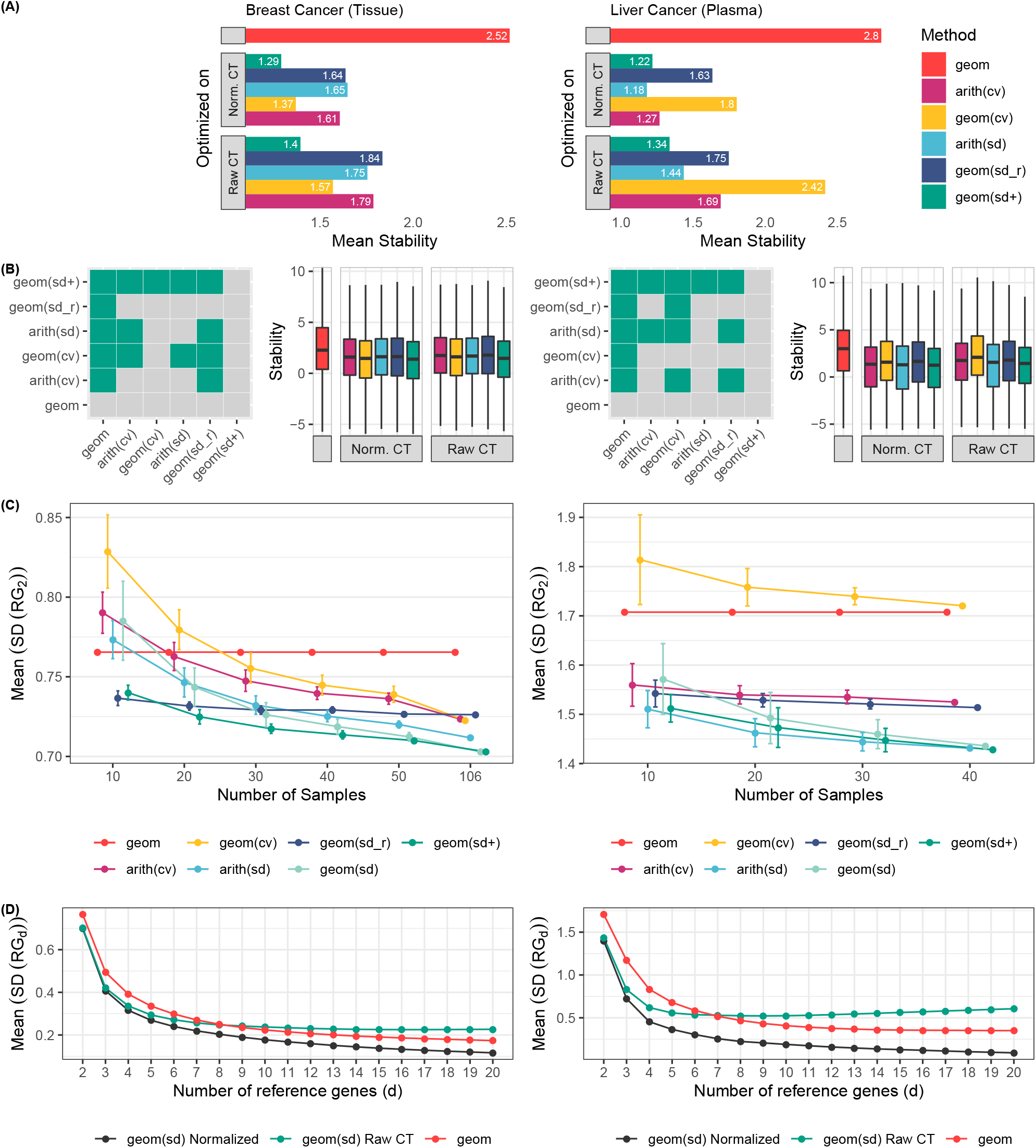
Stability comparison of different weighting methods. Figures on the left and right side are for the breast cancer and liver cancer respectively. (A) Mean stability of all combinations of two miRNAs in different weighting methods (the lower is better). (B) Box plots for the stability of all combinations of two miRNAs in different weighting methods and the tile figures show paired Wilcoxon test between the stability of different weighting methods on raw CT values. Colored tiles indicate that the row weighting method had significantly (p*<*0.01) lower stability than the column one. (C) Sample size analysis: for each sample size, the SD of each combination of two miRNAs is calculated and averaged. This process is repeated 20 times, and the error bars show the standard deviation of the repeats. The weights were calculated based on raw CT values. A statistical comparison in each sample size is provided in supplementary Figure S9 and S10 (D) The number of reference genes effect on stability. For each number of reference genes, the SD of different combinations of miRNAs were calculated. SD: Standard Deviation, Normalized: weights were optimized on the normalized data, raw CT: weights were optimized on the raw CT values, geom: usual geometric mean. Also for complete figures of all stability measures refer to the supplementary material

**Table 1.**
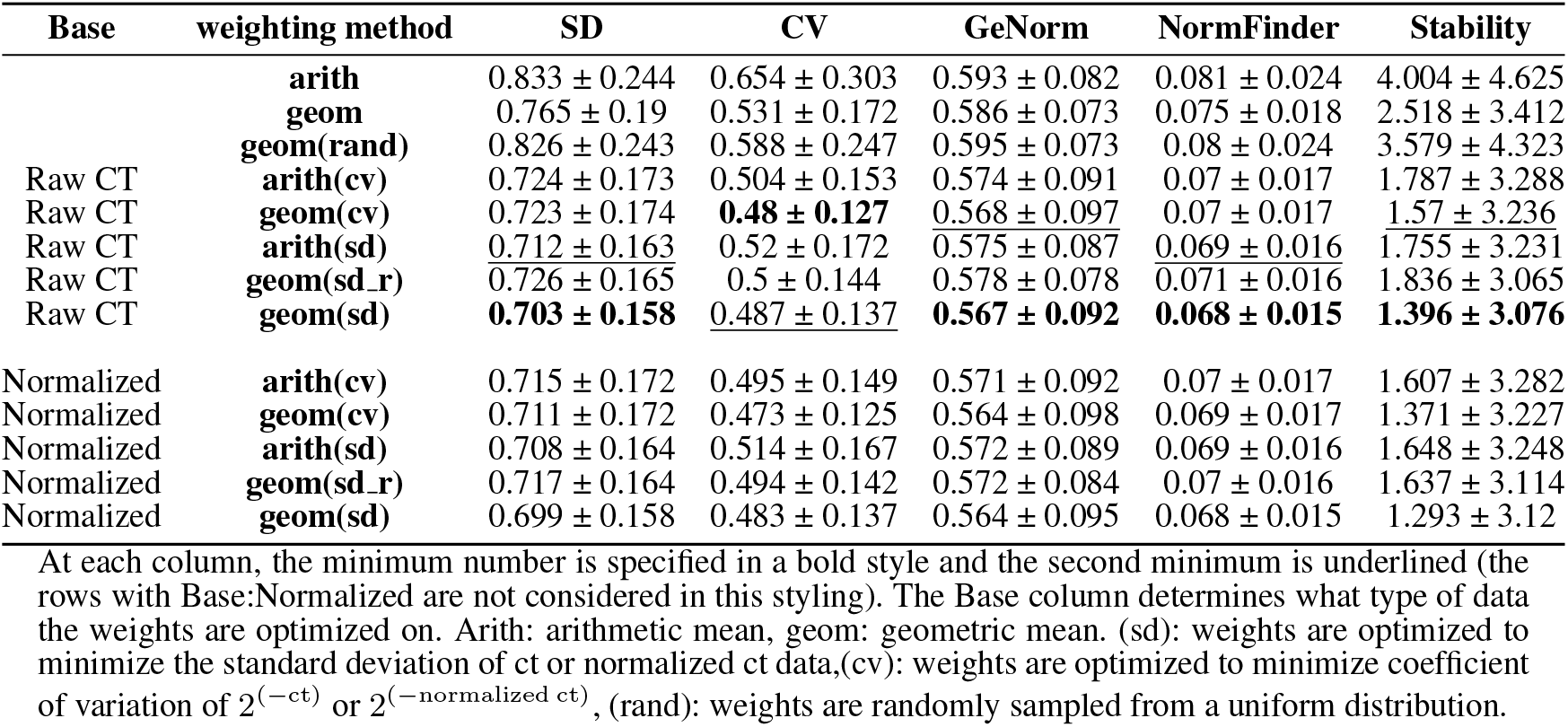
Comparison between different weighting methods based on four stability measures on the **breast cancer** dataset.

**Table 2.**
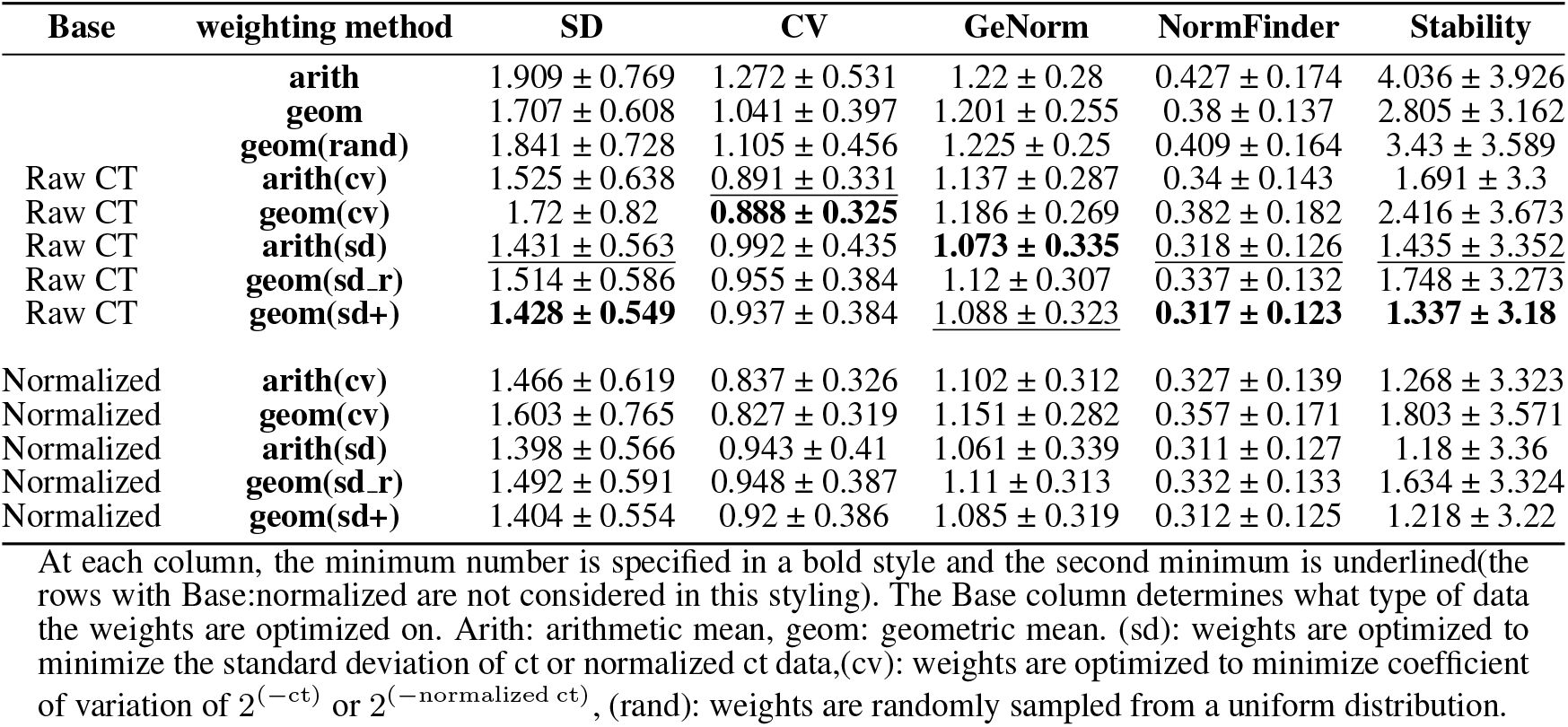
Comparison between different weighting methods based on four stability measures on the **liver cancer** dataset.

### 4.2 Sample Size Analysis

We analyzed the effect of sample size on the performance of the weighting methods. Each sub-sampling is repeated 20 times, and the mean standard deviation of all RG_2_ combinations are presented. The weights were calculated from the raw CT values and confidence intervals are provided. results are presented in Fig.2C. As expected, increasing the sample size improves all methods. Specifically geom(sd+), the improved version of geom(sd) for small sample sizes, shows significantly better results in lower samples sizes and As the number of samples decrease the improvement gap between geom(sd+) and geom(sd) increases. On the breast cancer dataset, the only methods which outperformed the usual geometric mean for the low sample size of 10 were geom(sd+) and geom(sd̲r) and as samples increased, although geom(sd+) kept improving, geom(sd̲r) stayed still. On the liver cancer dataset, all methods except geom(cv) performed better than the usual geometric mean independent of the sample size.

### 4.3 Number of Reference Genes

In order to figure out how the number of RGs affects the geom(sd) method, an iterative approach was used. Starting with *k* = 2 first, we evaluated all combinations of RG_2_ in terms of standard deviation. Then at each iteration *k*, the top 400 combinations of RG_k_ with the lowest SD were crossed with the remaining genes to build the combinations of RG_k+1_. This process was repeated till the number of RGs reached 20. Fig.2D shows the relation between the number of RGs (k) and the mean standard deviation of RG_k_ combinations. Until *k* = 6 the geom(sd) weighting method outperforms geom. However, when the weights are calculated based on the raw CT values, an over-fitting pattern appears. This result suggests that the geom(sd) weighting method is only applicable to up to five or six RGs.

### 4.4 Weights from External Dataset

Here we analyse whether an external high-throughput data could be used to calculate the aggregation weights of RGs. The TCGA breast cancer miRNA expression dataset was obtained as a biologically compatible external dataset for the qPCR array breast cancer dataset. Moreover, the qPCR array dataset is evaluated in three different sample sizes, 20, 30, and 106, just like what we saw in Fig2 geom(sd+) outperformed other methods. The paired Wilcoxon test reveals that calculating geom(sd+) weights from the miRNA-Seq data had better stability results compared to the raw CT with 20 samples but performed on par with the 30-sample case (Fig3C). On the other hand, the arith(sd) and arith(cv) methods were utterly off by a large margin regarding stability for the miRNA-Seq case.

**Fig. 3:**
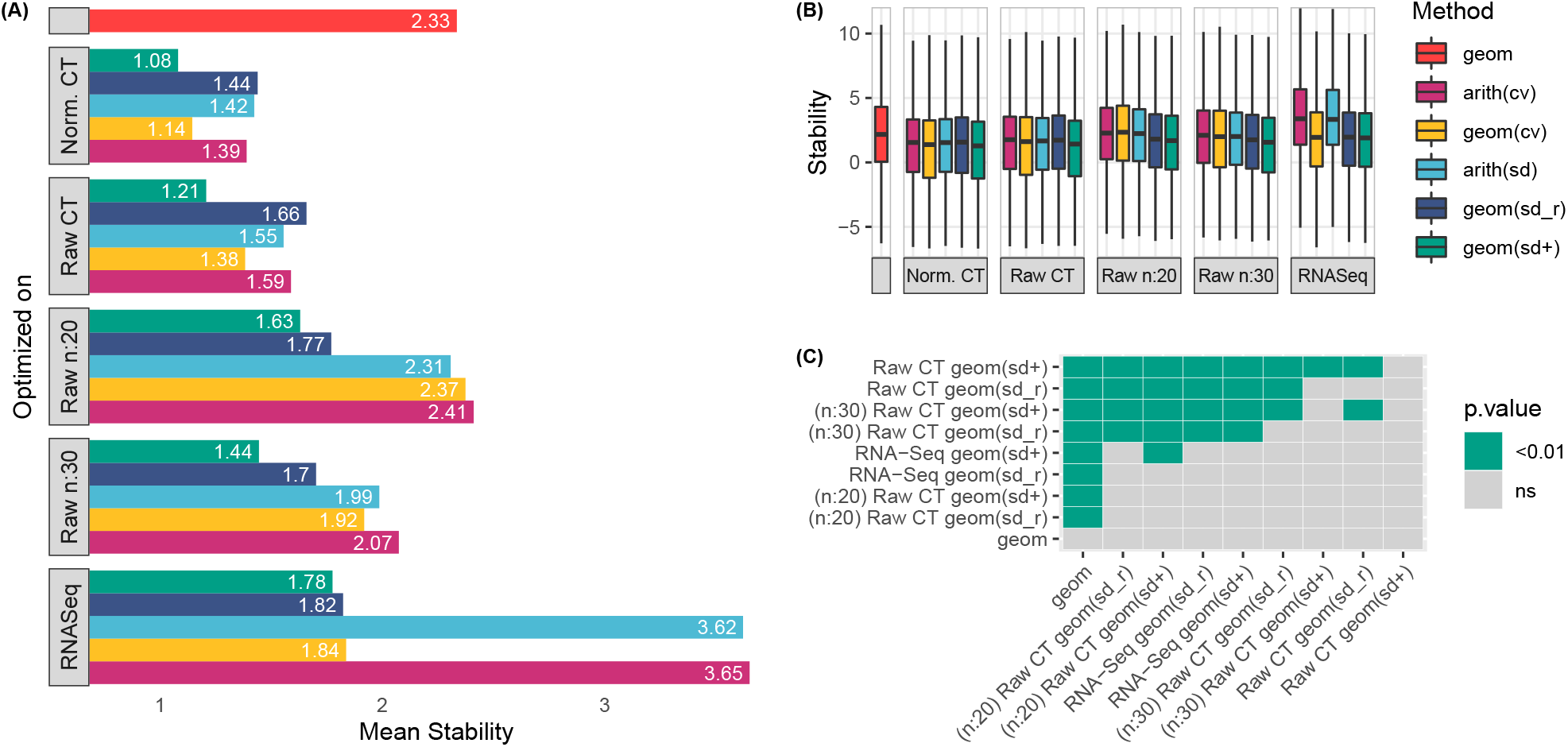
Comparison between external RNA-seq data and raw CT qPCR data with different sample sizes for weights optimization. The estrogen positive samples of the TCGA BRCA are used as the external RNA-seq dataset. (A) Mean stability of all combinations of two miRNAs in different weighting methods. (B) Box plots for the stability of all combinations of two miRNAs in different weighting methods. (C) Paired Wilcoxon test between stability scores of different weighting methods. Cells with *p <* 0.01 indicate that the row’s weighting method had significantly lower stability scores than the column one (lower is better). Raw CT: weights were optimized on the raw CT values of the entire 106 sample breast cancer qPCR array. n:x means a subset of x samples was taken and an average score of repeating the sub-sampling 20 times was considered.

### 4.5 Experimental Validation

To demonstrate the utility of the weighting method in a real experiment, we applied it to qPCR data of breast cancer tissue containing expressions of 3 internal controls (miR-16-5p, miR-361-5p, and RNU48) and one target miRNA (miR-21-5p). Fig4 illustrates how normalizing the expression of the well known miRNA miR-21-5p using a weighted geometric mean of the internal controls yielded a significant differential expression. In contrast, the usual geometric mean showed no significant change.

**Fig. 4:**
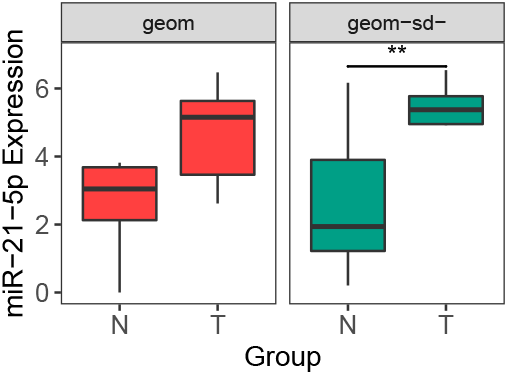
hsa-miR-21-5p expression normalized with the proposed weight method geom(sd) and normal geometric mean of three internal controls: U48, hsa-miR-16-5p and hsa-miR-361-5p. **: *t*-test p-value *<* 0.01

## 5 DISCUSSION

In this paper the aggregation of multiple RGs is introduced as an optimization problem. This optimization is formulated in four combinations of stability measures (SD or CV) as the objective functions and weighted mean (geometric or arithmetic) as aggregation functions. The geom(sd) method showed significantly better results compared to other methods in both low and high variability conditions as well as different numbers of samples. We also mathematically showed that weights of geom(sd) method are independent of the noise caused by the RNA abundance in different samples, which may justify its superiority over other methods. Furthermore, its closed form and regression-based solution allows fast running time and straightforward implementation in various platforms. We have also highlighted how a significant up-regulation of miR-21-5p could be overlooked in a real-world case if the non-weighted geometric mean was applied instead. miR-21-5p is a well-known up-regulated gene in cancer, as it is involved in cell growth and proliferation (Wang *et al*., 2019).

Through optimization of stability measures, our proposed aggregation methods provide either a more stable virtual RG or, at worst, an equal level of stability compared to the single input RGs. It is also noteworthy that the application of the usual geometric mean could result in a less stable virtual RG (supplementary material of Andersen *et al*. (2004)) specifically when there is a positive covariance between the RGs expression. the geom(sd r) method which uses the reverse of each RG’s SD as aggregation weights is also subject to this problem due to not considering the covariance between RGs (Qureshi and Sacan, 2013). In Eq.17 we demonstrated that in order to minimize the standard deviation of the aggregated virtual RG for a combination of two RGs, their covariance must be taken into account.

This study found that, when enough samples are available, raw CT values are preferable to external high throughput data for optimizing the aggregation weights of RGs. In our experiment, we used a compatible large sample size TCGA breast cancer miRNA-Seq data, yet a subset of 30 samples from the qPCR dataset showed better results. This can be a consequence of the distribution shift caused by the platform difference or batch effect. Considering these findings, in Fig.5 a workflow for the use cases of the InterOpt tool is presented. There are two main scenarios in which this tool would be useful. The first and most common scenario is when the RGs are preselected, or the experiment is already done. Then based on the number of samples and availability of a high-throughput data with similar biological conditions, the weights would be either calculated based on the raw CT values or the normalized high-throughput dataset. In the second scenario this tool can also be used to choose the best weighted combination of RGs from a qPCR experiment of common RGs or a high-throughput dataset. This use case is more suitable for situations where there is no consensus on the best combination of RGs for a particular biological condition. It is worth noting that to have a persistent and reliable result while using an external high-throughput dataset, the similarity of the clinical and pathological characteristics of samples with the qPCR study is highly recommended for both choosing the RGs or calculating the weights of the preselected RGs.

**Fig. 5:**
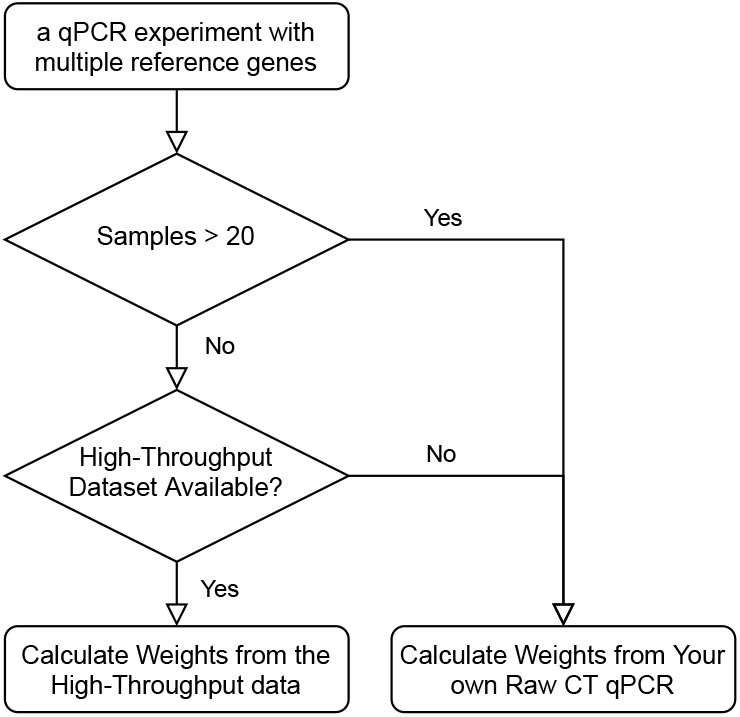
recommended usage of the proposed method

A limitation of the proposed method is the number of RGs. As described in section 4.3, aggregating more than 5 RGs by geom(sd) method doesn’t have the expected benefits compared to the regular geometric mean. But it is worth noting in most cases, no more than 3 RGs are quantified. Moreover, the number of samples for evaluating the hsa-miR-21-5p expression in the experimental validation phase was low (12 pairs) Due to the effect of gene expression distribution assumption on the RG stability measure, examining other distributions (like beta distribution) and providing better measures are suggested for further improvements. Moreover, we introduced the family of scale invariant functions as a necessary condition for aggregating multiple RGs. This family of functions can also be explored for more stable members in this line of research.

## ACKNOWLEDGEMENTS

Authors are thankful of Prof. Seyed Javad Mowla from Tarbiat Modares University for his contribution to this work. Also authors thank *greg*, a user from the math.stackexchange.com website for his assistance with some of mathematical solutions. Authors declare no conflicting interests or financial support.

