## Appendix for "InterOpt: Improved gene expression quantification in qPCR experiments using weighted aggregation of reference genes"

**Theorem 1 ( Criterion of optimum reference gene)**

A widely used application of gene expression quantification is differential expression analysis. In which genes expression are compared among samples using fold change or ratio:

$$I = \frac{y_{i,a}}{y_{i,b}} \tag{1}$$

Here $y_{i,a}$ is the expression of gene $i$ in sample $a$ and $y_{i,b}$ is the expression of gene $i$ in sample $b$. We aim to find the biological variations however, the CT values of a qPCR experiment also comprise technical variations. One of the primary sources of technical variation is the different amounts of initial RNA concentration at the start of the qPCR process for each sample. We can model this effect as a coefficient for each sample's different genes:

$$I' = \frac{\alpha_a y_{i,a}}{\alpha_b y_{i,b}} \tag{2}$$

Here $\alpha_a y_{i,a}$ and $\alpha_b y_{i,b}$ are the raw measured concentration of gene $i$ in samples $a$ and $b$ accordingly and $\alpha_a$ and $\alpha_b$ represent the technical variation as coefficients. In order to remove this technical variation, each gene expression is divided by an RG ($y_r$) which is also affected by the technical variation:

$$\frac{\alpha_a y_{i,a}}{\alpha_b y_{i,b}} = \frac{\frac{\alpha_a y_{i,a}}{\alpha_a y_{r,a}}}{\frac{\alpha_b y_{i,b}}{\alpha_b y_{r,b}}} = \frac{y_{i,a}}{y_{i,b}} \frac{y_{r,b}}{y_{r,a}} \approx \frac{y_{i,a}}{y_{i,b}} \tag{3}$$

To find the true ratio of the target gene, the ratio of the RG expression in different samples should be close to 1. $Z$ is a continuous random variable with probability density function $f_Z(z;\theta)$ representing this ratio. Hence the objective can be defined as maximizing the probability density of $Z$ in the proximity of 1:

$$\arg\max_\theta f_Z(z=1;\theta) \quad \equiv \quad \arg\max_\theta \left\{ \lim_{\epsilon \to 0} \int_{1-\epsilon}^{1+\epsilon} f_Z(z;\theta) \mathrm{d}z \right\} \tag{4}$$

$\theta$ represents the parameters of the probability density function.

**Theorem 2 (Modeling gene expression by Gaussian distribution)**

One of the common distributions to model gene expression is the Gaussian distribution. If we model the expression of an RG $r$ by a Gaussian distribution with mean $\mu$ and standard deviation $\sigma$, the distribution of the ratio of the gene in two different samples would be Eq.6 [2].

$$\frac{y_{r,b} \sim \mathcal{N}(\mu, \sigma^2)}{y_{r,a} \sim \mathcal{N}(\mu, \sigma^2)} = Z \tag{5}$$

$$\begin{aligned} f_Z(z) =& \frac{\mu(z+1).e^{-\frac{\frac{\mu^2}{\sigma^2}(z-1)^2}{2(z^2+1)}}}{\sigma\sqrt{2\pi}(\sqrt{z^2+1})^3}. \\ & \left[\Phi\left(\frac{\mu(z+1)}{\sigma\sqrt{z^2+1}}\right) - \Phi\left(-\frac{\mu(z+1)}{\sigma\sqrt{z^2+1}}\right)\right] + \frac{1}{(z^2+1)\pi} e^{-\frac{\mu^2}{\sigma^2}} \end{aligned} \tag{6}$$

$$a(z) = \sqrt{\frac{1}{\sigma_x^2} z^2 + \frac{1}{\sigma_y^2}},\ b(z) = \frac{\mu_x}{\sigma_x^2} z + \frac{\mu_y}{\sigma_y^2},\ d(z) = e^{\frac{b^2(z) - c a^2(z)}{2a^2(z)}},\ c = \frac{\mu_x^2}{\sigma_x^2} + \frac{\mu_y^2}{\sigma_y^2} \tag{7}$$

Considering a reference gene, we can assume that its distribution is the same over samples. Therefore we can rewrite it as follows:

$$a(z) = \frac{1}{\sigma} \sqrt{z^2 + 1},\ b(z) = \frac{\mu}{\sigma^2}(z+1),\ c = 2\frac{\mu^2}{\sigma^2} \tag{8}$$

$$d(z) = e^{\frac{\frac{\mu^2}{\sigma^4}(z+1)^2 - 2\frac{\mu^2}{\sigma^2}\frac{1}{\sigma^2}(z^2+1)}{2\frac{1}{\sigma^2}(z^2+1)}} \tag{9}$$

$$= e^{\frac{\frac{\mu^2}{\sigma^4}(z^2+1+2z) - 2\frac{\mu^2}{\sigma^2}\frac{1}{\sigma^2}(z^2+1)}{2\frac{1}{\sigma^2}(z^2+1)}} \tag{10}$$

$$= e^{\frac{\frac{\mu^2}{\sigma^2}(-z^2-1+2z)}{2(z^2+1)}} \tag{11}$$

$$= e^{-\frac{\frac{\mu^2}{\sigma^2}(z-1)^2}{2(z^2+1)}} \tag{12}$$

By plugging them into the Eq.6 we would have:

$$f_Z(z) = \frac{\frac{\mu}{\sigma^2}(z+1).e^{-\frac{\frac{\mu^2}{\sigma^2}(z-1)^2}{2(z^2+1)}}}{(\frac{1}{\sigma}\sqrt{z^2+1})^3}.\frac{1}{\sqrt{2\pi}\sigma^2}\left[\Phi\left(\frac{b(z)}{a(z)}\right) - \Phi\left(-\frac{b(z)}{a(z)}\right)\right] + \frac{1}{(z^2+1)\pi} e^{-\frac{\mu^2}{\sigma^2}} \tag{13}$$

$$= \frac{\mu(z+1).e^{-\frac{\frac{\mu^2}{\sigma^2}(z-1)^2}{2(z^2+1)}}}{\sigma\sqrt{2\pi}(\sqrt{z^2+1})^3}.\left[\Phi\left(\frac{\mu(z+1)}{\sigma\sqrt{z^2+1}}\right) - \Phi\left(-\frac{\mu(z+1)}{\sigma\sqrt{z^2+1}}\right)\right] + \frac{1}{(z^2+1)\pi} e^{-\frac{\mu^2}{\sigma^2}} \tag{14}$$

According to Eq.4 the goal is to maximize $f_z$ in $z = 1$. Then $f_z$ can be rewritten as a function of $k = \frac{\mu}{\sigma}$: .

$$k = \frac{\mu}{\sigma}, z = 1 \longrightarrow f_z(z=1,k) = \frac{k}{2\pi\sqrt{2}}.\left[\Phi\left(\sqrt{2}k\right) - \Phi\left(-\sqrt{2}k\right)\right] + \frac{1}{2\pi} e^{-k^2} \tag{15}$$

$$= \frac{k}{2\pi\sqrt{2}} \int_0^{k\sqrt{2}} 2e^{-\frac{t^2}{2}}\,\mathrm{d}t + \frac{1}{2\pi} e^{-k^2} \tag{16}$$

Now we can show that $\frac{\partial f_z(z=1,k)}{\partial k}$ is always a positive value:

$$\frac{\partial f_z(z=1,k)}{\partial k} = \frac{1}{2\sqrt{2}\pi} \int_0^{2k\sqrt{2}} e^{-\frac{t^2}{2}}\,\mathrm{d}t + \frac{2\sqrt{2}k}{2\sqrt{2}\pi} e^{-k^2} - \frac{2}{2\pi} k e^{-k^2} \tag{17}$$

$$= \frac{1}{2\sqrt{2}\pi} \int_0^{2k\sqrt{2}} e^{-\frac{t^2}{2}}\,\mathrm{d}t \geq 0 \tag{18}$$

Considering the Eq.4, it can be proven that maximization of $\frac{\mu}{\sigma}$ is equivalent to maximizing $f_Z(z = 1; \mu, \sigma)$ or in other words minimizing CV of the Gaussian distribution. This implies CV as a stability measure if the distribution of the RG expression follows the Gaussian distribution.

**Theorem 3 (Modeling gene expression by log-normal distribution)**

Another previously suggested distribution to model gene expression is log-normal distribution [1]. As the expression of a gene is always a positive number, this distribution has some benefits compared to the Gaussian distribution.

$$\log(X), \log(Y) \sim \mathcal{N}(\mu, \sigma^2) \tag{19}$$

$$Z' = \log(Z) = \log(X) - \log(Y) \longrightarrow Z' \sim \mathcal{N}(0, 2\sigma^2) \tag{20}$$

Now we can rewrite Eq.4 in terms of $Z'$:

$$\arg\max_{\theta} \int_{-\epsilon}^{\epsilon} P_{Z'}(z', \theta) \mathrm{d}z' = \arg\max_{\sigma} \int_{-\epsilon}^{+\epsilon} \mathcal{N}(0, 2\sigma^2) = \arg\min_{\sigma} \sigma \tag{21}$$

In conclusion, assuming gene expression follows a log-normal distribution, an RG with lower SD of the logarithm of expression is more stable.

**In previous sections we explained why standard deviation of log and coefficient of variation can be used as stability criteria. The next three theorems show how we can minimize those criteria in the context of weighted geometric/arithmetic mean of multiple reference genes.**

**Theorem 4 (Optimal weights for geometric mean to minimize SD of log expression)** *geom(sd)*

This section determines the optimal weighted geometric mean to minimize the SD of the logarithm of the aggregated RG. The geometric mean is equivalent to the arithmetic mean in the logarithmic space; therefore the optimization problem would be as follows:

$$\begin{gathered} \arg\min_{w_1, w_2, \ldots, w_d} \mathrm{SD}(\log(\prod_{i=1}^{d} y_{i,j}^{w_i})) \\ \text{subject to} \sum_{i=1}^{d} w_i = 1 \end{gathered} \tag{22}$$

$d$ is the number of reference genes and $y_{i,j}$ is the expression of gene $i$ in sample $j$. By applying logarithm, production converts to summation and Eq.22 can be rewritten as follows:

$$x_{i,j} = \log(y_{i,j}) \implies \mathrm{SD}(\sum_{i=1}^{d} w_i x_{i,j}) = \sum_{j=1}^{n}((\sum_{i=1}^{d} w_i x_{i,j}) - \frac{1}{n}(\sum_{k=1}^{n}(\sum_{i=1}^{d} w_i x_{i,k})))^2 \tag{23}$$

$$= \sum_{j=1}^{n}((\sum_{i=1}^{d} w_i x_{i,j}) - (\sum_{i=1}^{d} w_i \sum_{k=1}^{n}(\frac{x_{i,k}}{n})))^2 \tag{24}$$

$$= \sum_{j=1}^{n}(\sum_{i=1}^{d} w_i(x_{i,j} - \sum_{k=1}^{n}(\frac{x_{i,k}}{n})))^2 \tag{25}$$

$$= \sum_{j=1}^{n}(\sum_{i=1}^{d} w_i \tilde{x}_{i,j})^2 \tag{26}$$

Here $\tilde{x}$ is the mean centered version of $x$. Eq.26 can be interpreted as the cost function of classic linear regression with MSE cost function:

$$\arg\min_{w_1, w_2, \ldots, w_d} \sum_{j=1}^{n}(\sum_{i=1}^{d} w_i \tilde{x}_{i,j})^2 \equiv \sum_{i=1}^{d} w_i \tilde{X}_i = 0, \tag{27}$$

$$\text{subject to} \sum_{i=1}^{d} w_i = 1 \tag{28}$$

To get rid of constraints of $w_i$, we can rewrite the Eq.27 as Eq.32:

$$\begin{aligned}\sum_{i=1}^{d} w_i\tilde{X}_i &= \sum_{i=1}^{d-1} w_i\tilde{X}_i + w_d\tilde{X}_d && (29)\\ &= \sum_{i=1}^{d-1} w_i\tilde{X}_i + (1-\sum_{i=1}^{d-1} w_i)\tilde{X}_d && (30)\\ &= \sum_{i=1}^{d-1} w_i(\tilde{X}_i - \tilde{X}_d) + \tilde{X}_d && (31)\end{aligned}$$

$$\sum_{i=1}^{d-1} w_i(\tilde{X}_d - \tilde{X}_i) - \tilde{X}_d = 0 \quad \equiv \quad \sum_{i=1}^{d-1} w_i G_i = \tilde{X}_d \tag{32}$$

Here $\tilde{X}_i$ is the mean centered expression of reference gene i and $G_i = \tilde{X}_d - \tilde{X}_i$. The solution of Eq.32 linear regression comes in closed form:

$$\begin{gathered} W_{1..(d-1)} = (G^T G)^{-1} G^T \tilde{X}_d, \quad w_d = 1 - \sum_{i=1}^{d-1} w_i \\ G = \begin{bmatrix} \tilde{X}_d - \tilde{X}_1 \\ \tilde{X}_d - \tilde{X}_2 \\ \vdots \\ \tilde{X}_d - \tilde{X}_{d-1} \end{bmatrix} \end{gathered} \tag{33}$$

$W_{1..(d-1)}$ is a $1 \times (d-1)$ matrix consisting of the first $(d-1)$ elements of $W$.

**Theorem 5 (Optimal weights for arithmetic mean to minimize CV)** *arith(cv)*

Suppose $W$ is a $1 \times d$ matrix and $M$ is a $d \times n$ matrix containing the expression of $d$ reference genes in $n$ samples. Here the goal is to minimize the CV of the weighted arithmetic mean of the reference genes. This optimization problem is demonstrated in Eq.34:

$$\begin{gathered} \arg\min_W \frac{\text{SD}(WM)}{\text{Mean}(WM)} \\ \sum_{i=1}^{d} w_i = 1 \end{gathered} \tag{34}$$

Eq.34 can be written as:

$$\begin{gathered} \arg\min_W \frac{\sqrt{\frac{1}{n}\left\|WM - \frac{1}{n}WM\mathbb{1}_n\right\|_2^2}}{\frac{1}{n}WM\mathbb{1}_n} \\ \sum_{i=1}^{d} w_i = 1 \end{gathered} \tag{35}$$

where $\mathbb{1}_n$ is a $n \times 1$ matrix of ones.
Introduce an unconstrained vector $x$ and use it to construct a column vector $w$, which satisfies the constraint.

$$w = \frac{x}{{\mathbb{1}_\mathtt{n}}^T x} \quad \Longrightarrow \quad \mathbb{1}^T w = \frac{\mathbb{1}^T x}{\mathbb{1}^T x} \doteq \mathbb{1} \tag{36}$$

Then for algebraic convenience, define some auxiliary variables

$$\mathbb{1} = A, \qquad J = AA^T \tag{37}$$

$$C = I - \tfrac{1}{n}J \qquad \text{(Centering Matrix)} \tag{38}$$

$$w = W^T \qquad \text{(column vector constructed from } x\text{)} \tag{39}$$

$$y = M^T w \qquad \implies dy = M^T dw \tag{40}$$

$$z = Cy \qquad \implies dz = CM^T dw \tag{41}$$

$$\alpha = \mathbb{1}^T x \qquad \implies d\alpha = \mathbb{1}^T dx \tag{42}$$

$$\beta = \mathbb{1}^T y \qquad \implies d\beta = \mathbb{1}^T dy = \mathbb{1}^T M^T dw \tag{43}$$

$$w = \alpha^{-1} x \qquad \implies dw = \alpha^{-1} dx - x\alpha^{-2} d\alpha \tag{44}$$

$$\implies dw = \alpha^{-1}(I - w\mathbb{1}^T)\, dx \tag{45}$$

Note that $C^T = C = C^2$ and $\beta = \mathbb{1}^T M^T w = w^T M \mathbb{1} = WMA$
these properties will be used in several of the steps below. Use the new variables to simplify the vector appearing in the numerator.

$$\left(WM - \tfrac{1}{n}WMAA^T\right)^T = \left(M^T w - \tfrac{1}{n}JM^T w\right) = Cy = z \tag{46}$$

Call the objective function $\phi$, and start by differentiating its square.

$$\phi^2 = n\beta^{-2} z^T z \tag{47}$$

$$2\phi\, d\phi = 2n\beta^{-2} z^T dz - 2n\beta^{-3} z^T z\, d\beta \tag{48}$$

$$d\phi = n\phi^{-1}\beta^{-3} z^T \left(\beta\, dz - z\, d\beta\right) \tag{49}$$

$$= n\phi^{-1}\beta^{-3} z^T \left(\beta\, CM^T - z\mathbb{1}^T M^T\right) dw \tag{50}$$

$$= n\phi^{-1}\alpha^{-1}\beta^{-3} z^T \left(\beta\, CM^T - z\mathbb{1}^T M^T\right)\left(I - w\mathbb{1}^T\right) dx \tag{51}$$

$$\frac{\partial\phi}{\partial x} = n\phi^{-1}\alpha^{-1}\beta^{-3}\left(I - \mathbb{1}w^T\right)\left(\beta\, MC - M\mathbb{1}z^T\right) z \tag{52}$$

Set the gradient to zero.

$$\left(\mathbb{1}w^T\right)\left(\beta\, MC - M\mathbb{1}z^T\right) z = I\left(\beta\, MC - M\mathbb{1}z^T\right) z \tag{53}$$

Eliminate $z$ in favor of $w$.

$$\left(\mathbb{1}w^T\right)\left(\beta MC - M\mathbb{1}w^T MC\right) CM^T w = \left(\beta MC - M\mathbb{1}w^T MC\right) CM^T w \tag{54}$$

$$\left(\mathbb{1}w^T\right)\left(\beta I - M\mathbb{1}w^T\right) MCM^T w = \left(\beta I - M\mathbb{1}w^T\right) MCM^T w \tag{55}$$

$$\left(\beta\mathbb{1}w^T - \mathbb{1}w^T M\mathbb{1}w^T\right) MCM^T w = \left(\beta I - M\mathbb{1}w^T\right) MCM^T w \tag{56}$$

$$0 = \left(\beta I - M\mathbb{1}w^T\right) MCM^T w \tag{57}$$

$$M\mathbb{1}w^T \sigma MCM^T w = \beta\, \sigma MCM^T w \tag{58}$$

$$\left((M\mathbb{1})w^T\right)\sigma v = \beta\sigma v \tag{59}$$

$$Bv = \beta v \tag{60}$$

The last line is an eigenvalue equation. Since the matrix $B$ is rank-$\mathbb{1}$, there is only one non-trivial eigenvector, which miraculously allows for a closed-form solution to the problem.

$$v = M\mathbb{1} \qquad \text{(eigenvector of } B\text{)} \tag{61}$$

$$(MCM^T)w = M\mathbb{1} \tag{62}$$

$$w = (MCM^T)^+ M\mathbb{1} + \left(I - (MCM^T)^+ MCM^T\right) q \tag{63}$$

where $H^+$ denotes the pseudo-inverse of $H$ and $q$ is an arbitrary vector.

**Theorem 6 (Optimal weights for geometric mean to minimize CV)** *geom(cv)*

This theorem is another variation of the previous one. The difference here is that instead of arithmetic mean, we have geometric mean. To write geometric mean as a matrix product here, we use an alternative form $\exp(W\ln(M))$. so if $M' = \ln(M)$, then the optimization problem would become:

$$\begin{gathered}\arg\min_{W} \frac{\text{SD}(\exp(WM'))}{\text{Mean}(\exp(WM'))} \\ \sum_{i=1}^{d} w_i = 1\end{gathered} \tag{64}$$

also Eq.64 can be written as:

$$\arg\min_{W} \frac{\sqrt{\frac{1}{n}\left\|\exp(WM') - \frac{1}{n}\exp(WM')A\right\|_2^2}}{\frac{1}{n}\exp(WM')A}, \quad \sum_{i=1}^{d} w_i = 1 \tag{65}$$

A is a $n \times 1$ matrix of ones.
Introduce an unconstrained vector $x$ and use it to construct a column vector $w$ which satisfies the constraint.

$$w = \frac{x}{\mathbb{1}^T x} \quad \implies \quad \mathbb{1}^T w = \frac{\mathbb{1}^T x}{\mathbb{1}^T x} \doteq \mathbb{1} \tag{66}$$

Then for algebraic convenience, define some auxiliary variables

$$\begin{aligned}
&\mathbb{1} = A, && J = AA^T && (67)\\
&C = I - \tfrac{1}{n}J && (\text{Centering Matrix}) && (68)\\
&w = W^T && (\text{column vector constructed from } x) && (69)\\
&B = \text{d×1 matrix of ones} && && (70)\\
&Q = M' \circ By^T && \circ : \text{Hadamard Product} && (71)\\
&y = exp(M'^T w) && \implies dy = M'^T \circ exp(M'^T w)B^T dw = Q^T dw && (72)\\
&z = Cy && \implies dz = CQ^T dw && (73)\\
&\alpha = \mathbb{1}^T x && \implies d\alpha = \mathbb{1}^T dx && (74)\\
&\beta = \mathbb{1}^T y && \implies d\beta = \mathbb{1}^T dy = \mathbb{1}^T Q^T dw && (75)\\
&w = \alpha^{-1} x && \implies dw = \alpha^{-1} dx - x\alpha^{-2} d\alpha && (76)\\
& && \implies dw = \alpha^{-1}(I - w\mathbb{1}^T)\, dx && (77)
\end{aligned}$$

Note that $C^T = C = C^2$ and $\beta = \mathbb{1}^T \exp(M'^T w) = \exp(w^T M')\mathbb{1} = \exp(WM')A$
these properties will be used in several of the steps below.
Use the new variables to simplify the vector appearing in the numerator.

$$\left(\exp(WM') - \tfrac{1}{n}\exp(WM')AA^T\right)^T = \left(\exp(M'^T w) - \tfrac{1}{n}J\exp(M'^T w)\right) = Cy = z \tag{78}$$

Call the objective function $\phi$, and start by differentiating its square.

$$\phi^2 = n\beta^{-2} z^T z \tag{79}$$

$$2\phi\, d\phi = 2n\beta^{-2} z^T dz - 2n\beta^{-3} z^T z\, d\beta \tag{80}$$

$$d\phi = n\phi^{-1}\beta^{-3} z^T \left(\beta\, dz - z\, d\beta\right) \tag{81}$$

$$= n\phi^{-1}\beta^{-3} z^T \left(\beta\, CQ^T - z\mathbb{1}^T Q^T\right) dw \tag{82}$$

$$= n\phi^{-1}\alpha^{-1}\beta^{-3} z^T \left(\beta\, CQ^T - z\mathbb{1}^T Q^T\right)\left(I - w\mathbb{1}^T\right) dx \tag{83}$$

$$\frac{\partial\phi}{\partial x} = n\phi^{-1}\alpha^{-1}\beta^{-3}\left(I - \mathbb{1}w^T\right)\left(\beta\, QC - Q\mathbb{1}z^T\right) z \tag{84}$$

Set the gradient to zero.

$$\left(\mathbb{1}w^T\right)\left(\beta\,QC - Q\mathbb{1}z^T\right)z = I\left(\beta\,QC - Q\mathbb{1}z^T\right)z \tag{85}$$

$$\tag{86}$$

Define the matrix

$$H = \beta QC - Q\mathbb{1}z^T \tag{87}$$

Then the zero gradient condition becomes

$$Hz = \left(\mathbb{1}w^T\right)Hz \;=\; \left(w^T Hz\right)\mathbb{1} \;=\; \sigma\mathbb{1} \tag{88}$$

This eliminates $z$ from one side of the equation

$$\left(\beta QC - Q\mathbb{1}z^T\right)z = \sigma\mathbb{1} \tag{89}$$

Other substitutions can be used

$$\beta = \mathbb{1}^T y \tag{90}$$

$$Cz = C^2 y = Cy = z \tag{91}$$

$$Y = \operatorname{Diag}(y) \qquad\Longrightarrow\qquad Q = MY, \quad z = Cy = CY\mathbb{1} \tag{92}$$

to rewrite the equation entirely in terms of $y$ (or $Y$)

$$\sigma\mathbb{1}_d = M\operatorname{Diag}(y)\Big(\left(\mathbb{1}_n^T y\right)C\operatorname{Diag}(y) - \left(y^T Cy\right)I_n\Big)\mathbb{1}_n \tag{93}$$

$$= M\Big(\left(\mathbb{1}_n^T Y\mathbb{1}_n\right)YCY - \left(\mathbb{1}_n^T YCY\mathbb{1}_n\right)Y\Big)\mathbb{1}_n \tag{94}$$

$$= S\mathbb{1}_n \tag{95}$$

$$S\mathbb{1}_n = \sigma\mathbb{1}_d \quad\Longrightarrow\quad \frac{S\mathbb{1}_n}{\|S\mathbb{1}_n\|} = \frac{\mathbb{1}_d}{\sqrt{d}} \tag{96}$$

This looks promising. It almost looks like an eigenvalue equation; however $S$ is rectangular, so the $\mathbb{1}$ vectors on the RHS and LHS have different lengths. The next step is to come up with an iteration formula akin to the classic power iteration to solve for a $y$ vector (or $S$ matrix) such that the pseudo-eigenvalue equation is satisfied. Since a closed-form eigenvalue-like solution seems improbable, and since we already know how to calculate the gradient and the cost function for any value of $x,$ i.e.

$$\phi(x) = \left(n\beta^{-2}z^T z\right)^{1/2} \tag{97}$$

$$g(x) = n\phi^{-1}\alpha^{-1}\beta^{-3}\left(I - \mathbb{1}w^T\right)\left(\beta\,QC - Q\mathbb{1}z^T\right)z \tag{98}$$

A better idea is a numerical solution using a gradient-based method. The Barzilai-Borwein method is straightforward and effective. Initialize

$$x_0 = random \tag{99}$$

First step

$$g_0 = g(x_0) \tag{100}$$

$$x_1 = x_0 - \left(\frac{0.05\;\phi(x_0)}{g_0^T g_0}\right)g_0 \tag{101}$$

$$k = 1 \tag{102}$$

Subsequent steps

$$g_k = g(x_k) \tag{103}$$

$$x_{k+1} = x_k - \left(\frac{\left(x_k - x_{k-1}\right)^T\left(g_k - g_{k-1}\right)}{\left(g_k - g_{k-1}\right)^T\left(g_k - g_{k-1}\right)}\right)g_k \tag{104}$$

$$k = k + 1 \tag{105}$$

Stop when $g_k \approx 0.$

In Practice, the original Barzilai-Borwein method could diverge from the optimal point. To handle this inconvenience, the gradient cautiously is controlled using the stabilized Barzilai-Borwein method.

**Theorem 7 (Effect of raw qPCR data technical biases on geom(sd) method)**

We found a solution to minimize the SD of the weighted geometric mean of multiple reference genes in this theorem. Here, we solve this problem in the context of random variables. Then by modeling technical variation and applying it to the reference genes, we show that the technical variation would not affect the solution.
Suppose $X_1$ and $X_2$ are two random variables representing the logarithm of two reference genes expression. The variance of the weighted arithmetic mean of them would be:

$$\text{var}(\frac{w_1X_1+w_2X_2}{w_1+w_2}) = \frac{w_1^2}{(w_1+w_2)^2}\text{var}(X_1) + \frac{w_2^2}{(w_1+w_2)^2}\text{var}(X_2) + 2\frac{w_1\cdot w_2}{(w_1+w_2)^2}\text{cov}(X_1,X_2) \tag{106}$$

$w_1$ and $w_2$ are the weights of $X_1$ and $X_2$. To find the minimum of this equation, we set the derivative to zero with respect to each of the weights:

$$\begin{aligned}\frac{\text{d}\{\text{var}(\frac{w_1X_1+w_2X_2}{w_1+w_2})\}}{\text{d}w_1} = &\frac{2w_1(w_1+w_2)^2 - 2w_1^2(w_1+w_2)}{(w_1+w_2)^4}\text{var}(X_1)+\\ &\frac{-2w_2^2(w_1+w_2)}{(w_1+w_2)^4}\text{var}(X_2)+\\ &2\frac{w_2(w_1+w_2)^2 - 2w_1w_2(w_1+w_2)}{(w_1+w_2)^4}\text{cov}(X_1,X_2) = 0\end{aligned} \tag{107}$$

$$\begin{aligned}\frac{\text{d}\{\text{var}(\frac{w_1X_1+w_2X_2}{w_1+w_2})\}}{\text{d}w_1} = &(2w_1(w_1+w_2) - 2w_1^2)\text{var}(X_1) + (-2w_2^2)\text{var}(X_2)+\\ &2(w_2(w_1+w_2) - 2w_1w_2)\text{cov}(X_1,X_2) = 0\end{aligned} \tag{108}$$

After several steps of simplification, we have the Eq.109:

$$\begin{cases}\frac{\text{d}\{\text{var}(\frac{w_1X_1+w_2X_2}{w_1+w_2})\}}{\text{d}w_1} = 0 & \longrightarrow w_1w_2\text{var}(X_1) - w_2^2var(X_2) + (w_2^2 - w_1w_2)\text{cov}(X_1,X_2) = 0\\ \frac{\text{d}\{\text{var}(\frac{w_1X_1+w_2X_2}{w_1+w_2})\}}{\text{d}w_2} = 0 & \longrightarrow -w_1^2\text{var}(X_1) + w_1w_2\text{var}(X_2) + (w_1^2 - w_1w_2)\text{cov}(X_1,X_2) = 0\end{cases} \tag{109}$$

$$(w_1^2 + w_1w_2)\text{var}(X_1) + (-w_2^2 - w_1w_2)\text{var}(X_2) + (w_2^2 - w_1^2)\text{cov}(X_1,X_2) = 0 \tag{110}$$

$$w_1\text{var}(X_1) - w_2\text{var}(X_2) + (w_2 - w_1)\text{cov}(X_1,X_2) = 0 \tag{111}$$

$$\frac{\text{var}(X_2) - \text{cov}(X_1,X_2)}{\text{var}(X_1) - \text{cov}(X_1,X_2)} = \frac{w_1}{w_2} \tag{112}$$

Next if we assume that the sum of $w_1$ and $w_2$ is equal to 1 then the closed-form solution of $w_1$ and $w_2$ would be as follows:

$$\frac{\text{var}(X_2) - \text{cov}(X_1,X_2)}{\text{var}(X_1) - \text{cov}(X_1,X_2)} = \frac{w_1}{w_2}, \quad w_1 + w_2 = 1 \tag{113}$$

$$w_1 = \frac{\text{var}(X_2) - \text{cov}(X_1,X_2)}{\text{var}(X_1) + \text{var}(X_2) - 2\text{cov}(X_1,X_2)}, \quad w_2 = \frac{\text{var}(X_1) - \text{cov}(X_1,X_2)}{\text{var}(X_1) + \text{var}(X_2) - 2\text{cov}(X_1,X_2)} \tag{114}$$

These equations can also be rewritten this way:

$$\begin{aligned}p_1 = (\text{var}(X_1) - \text{cov}(X_1,X_2))^{-1}, \quad p_2 = (\text{var}(X_2) - \text{cov}(X_1,X_2))^{-1}\\ w_1 = \frac{p_1}{p_1+p_2}, \quad w_2 = \frac{p_2}{p_1+p_2}\end{aligned} \tag{115}$$

The expression values obtained from qPCR (CT values) are subject to technical and biological variations. Here we model the technical variation as an additive random variable called $F$. So we replace $X_1$ and $X_2$ with $X_1 + F$ and $X_2 + F$ in Eq.114. As $X_1$ and $X_2$ are in logarithmic space, adding $F$ to them is like applying a random coefficient to each of the samples' true expressions. Now we can simplify the equation as follows:

$$w_1 = \frac{\text{var}(X_2 + F) - \text{cov}(X_1 + F, X_2 + F)}{\text{var}(X_1 + F) + \text{var}(X_2 + F) - 2\text{cov}(X_1 + F, X_2 + F)} \tag{116}$$

$$= \frac{\text{var}(X_2) + \text{var}(F) - \text{cov}(X_1, X_2) - \text{var}(F)}{\text{var}(X_1) + \text{var}(F) + \text{var}(X_2) + \text{var}(F) - 2\text{cov}(X_1, X_2) - 2\text{var}(F)} \tag{117}$$

$$= \frac{\text{var}(X_2) - \text{cov}(X_1, X_2)}{\text{var}(X_1) + \text{var}(X_2) - 2\text{cov}(X_1, X_2)} \tag{118}$$

The same steps could be applied to $w_2$. As you can see, the result is the same as Eq.114. This suggests that if we assume the technical variation is independent of the gene and in the form of scale operations (additive in logarithmic space), the weights calculated from the raw CT values are not affected by technical variation.
