## Supplementary material for "InterOpt: Improved gene expression quantification in qPCR experiments using weighted aggregation of reference genes": Supplemetary

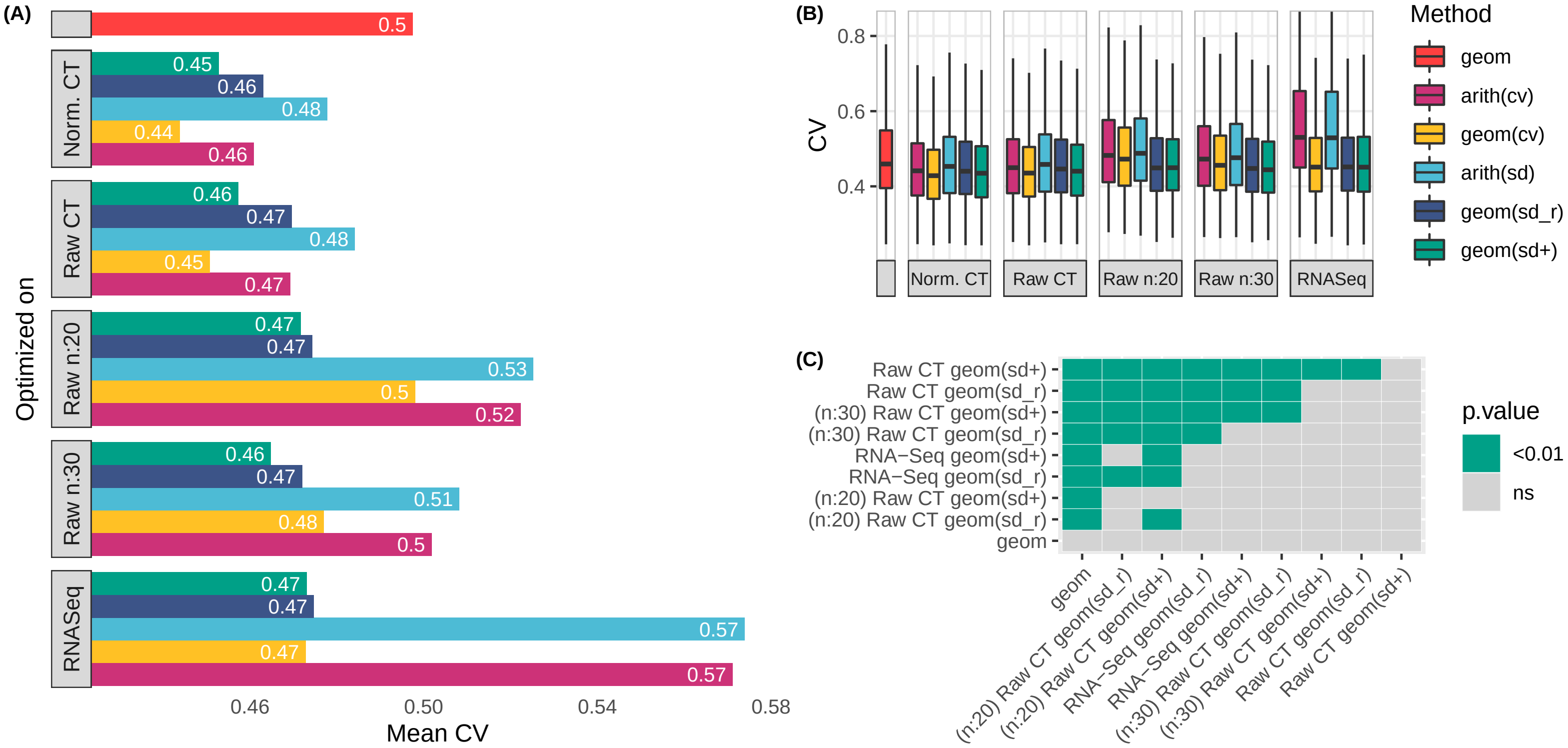


Figure S.1: Comparison between external RNA-seq data and raw CT qPCR data with different sample sizes for weights optimization. The estrogen positive samples of the TCGA BRCA are used as the external RNA-seq dataset. (A) Mean **CV** of all combinations of two miRNAs in different weight methods. (B) Box plots for the CV of all combinations of two miRNAs in different weight methods. (C) Paired Wilcoxon test between CV of different weight methods. Cells with $p < 0.01$ indicate that the row's weight method had significantly lower CV than the column one (lower is better). Raw CT: weights were optimized on the raw CT values of the entire 106 sample breast cancer qPCR array. n:x means a subset of x samples was taken and an average score of repeating the sub-sampling 20 times was considered.

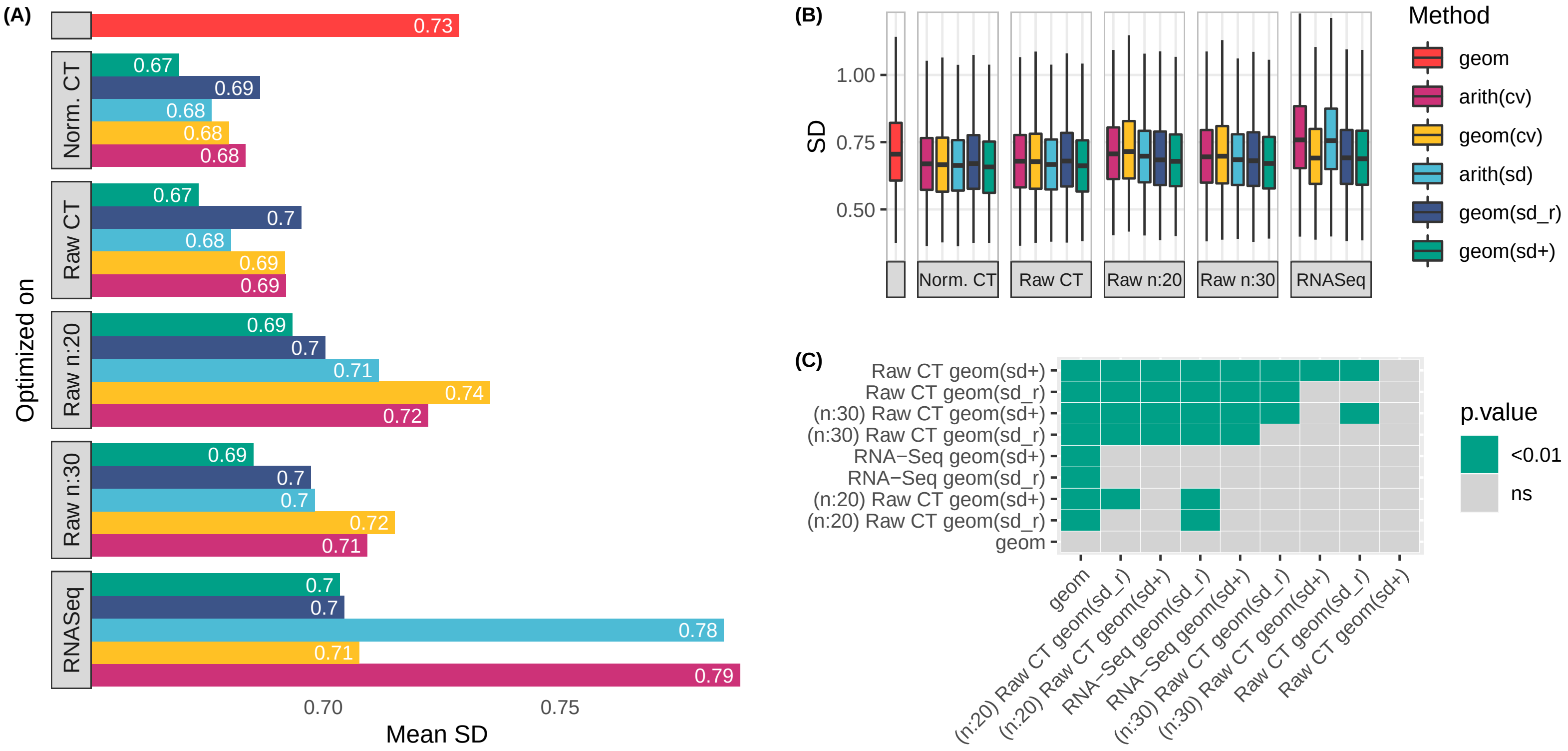


Figure S.2: Comparison between external RNA-seq data and raw CT qPCR data with different sample sizes for weights optimization. The estrogen positive samples of the TCGA BRCA are used as the external RNA-seq dataset. (A) Mean **SD** of all combinations of two miRNAs in different weight methods. (B) Box plots for the SD of all combinations of two miRNAs in different weight methods. (C) Paired Wilcoxon test between CV of different weight methods. Cells with $p < 0.01$ indicate that the row's weight method had significantly lower CV than the column one (lower is better). Raw CT: weights were optimized on the raw CT values of the entire 106 sample breast cancer qPCR array. n:x means a subset of x samples was taken and an average score of repeating the sub-sampling 20 times was considered.

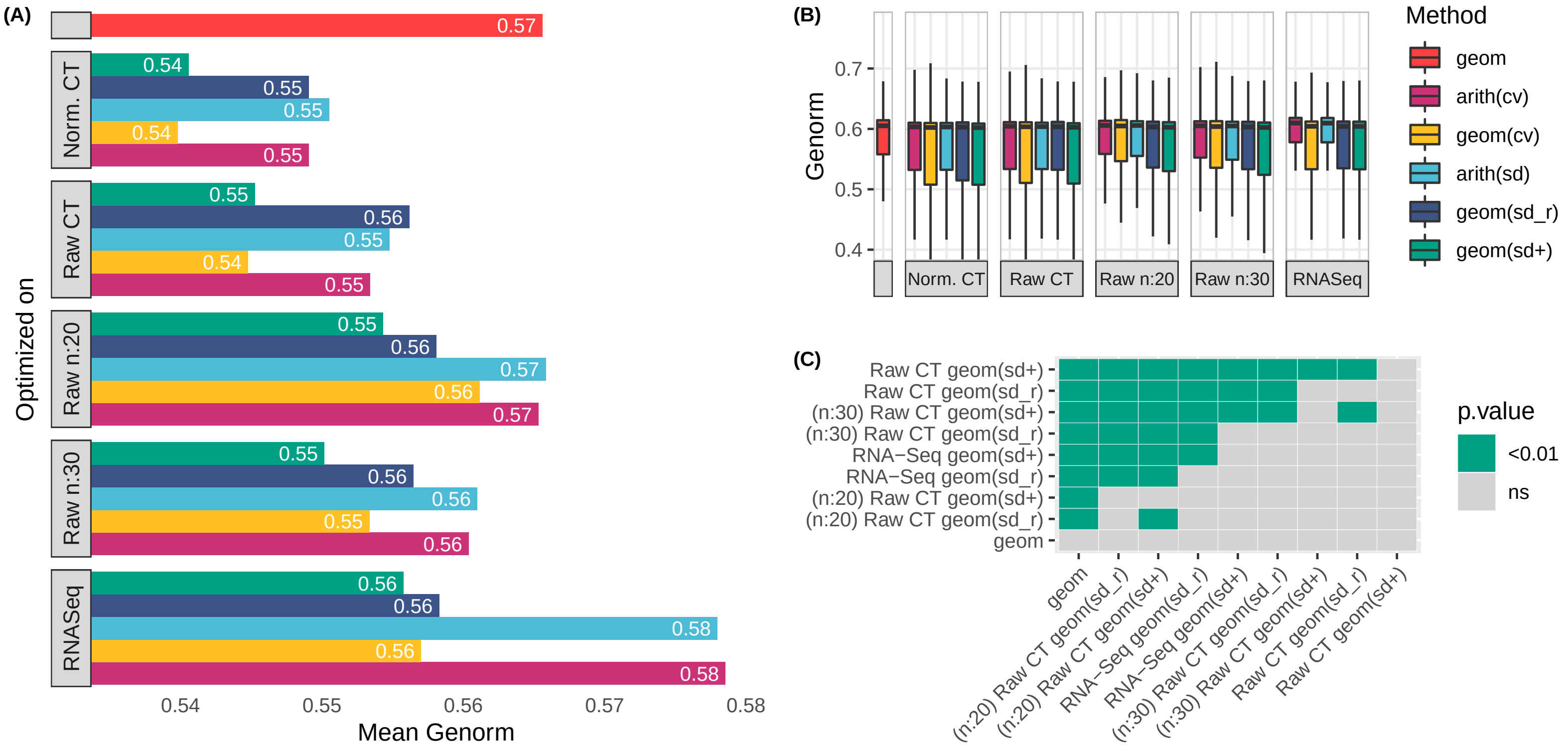


Figure S.3: Comparison between external RNA-seq data and raw CT qPCR data with different sample sizes for weights optimization. The estrogen positive samples of the TCGA BRCA are used as the external RNA-seq dataset. (A) Mean **Genorm** score of all combinations of two miRNAs in different weight methods. (B) Box plots for the Genorm score of all combinations of two miRNAs in different weight methods. (C) Paired Wilcoxon test between Genorm score of different weight methods. Cells with $p < 0.01$ indicate that the row's weight method had significantly lower Genorm score than the column one (lower is better). Raw CT: weights were optimized on the raw CT values of the entire 106 sample breast cancer qPCR array. n:x means a subset of x samples was taken and an average score of repeating the sub-sampling 20 times was considered.

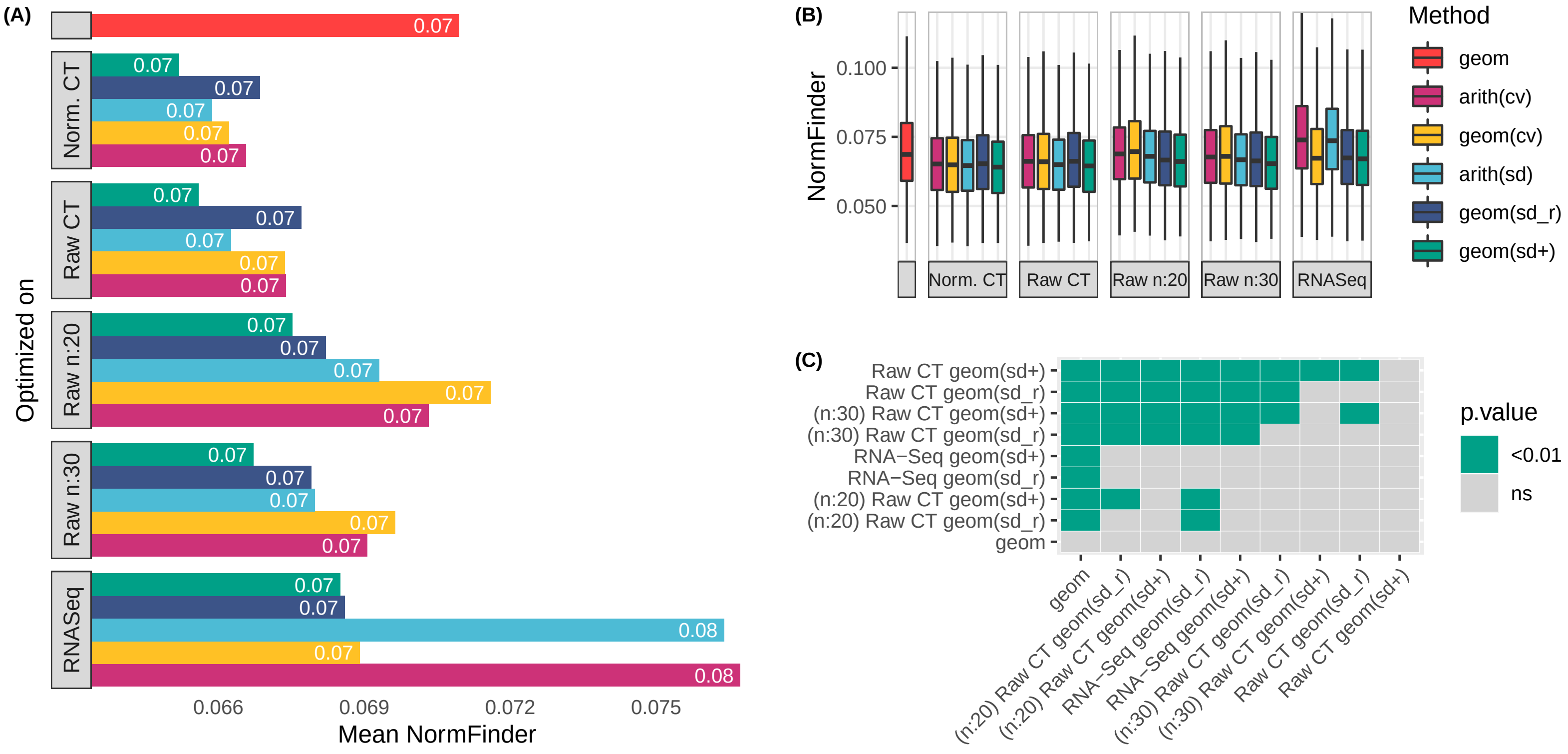


Figure S.4: Comparison between external RNA-seq data and raw CT qPCR data with different sample sizes for weights optimization. The estrogen positive samples of the TCGA BRCA are used as the external RNA-seq dataset. (A) Mean **NormFinder** score of all combinations of two miRNAs in different weight methods. (B) Box plots for the NormFinder score of all combinations of two miRNAs in different weight methods. (C) Paired Wilcoxon test between NormFinder score of different weight methods. Cells with $p < 0.01$ indicate that the row's weight method had significantly lower NormFinder score than the column one (lower is better). Raw CT: weights were optimized on the raw CT values of the entire 106 sample breast cancer qPCR array. n:x means a subset of x samples was taken and an average score of repeating the sub-sampling 20 times was considered.

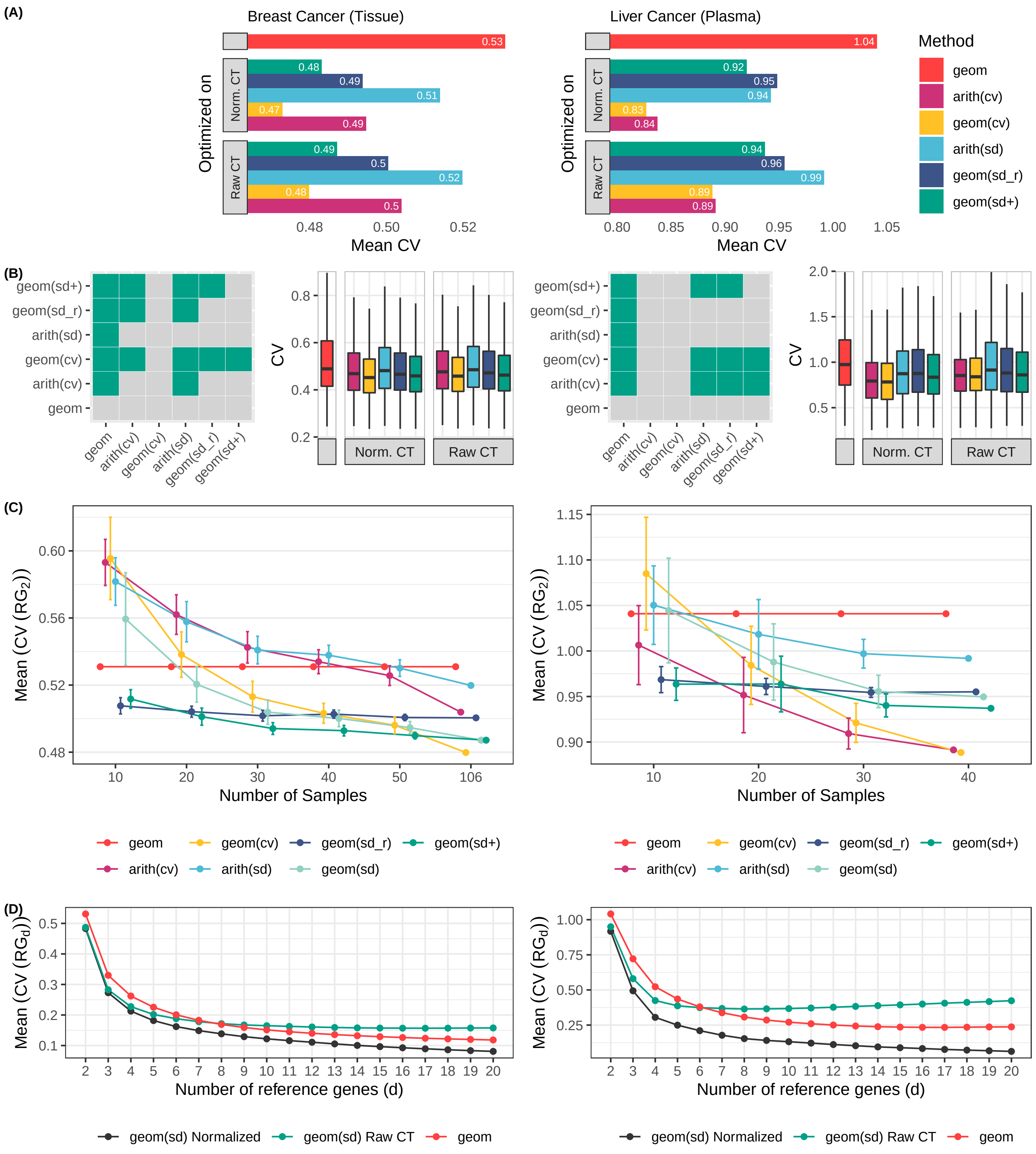


Figure S.5: Stability (**CV**) comparison of different weight methods. Figures on the left and right side are for the breast cancer and liver cancer respectively. (A) Mean stability of all combinations of two miRNAs in different weight methods (the lower is better). (B) Box plots for the stability of all combinations of two miRNAs in different weight methods and the tile figures show paired Wilcoxon test between the stability of different weight methods on Raw CT values. Colored tiles indicate that the row weight method had significantly ($p<0.01$) lower stability than the column one. (C) Sample size analysis: for each sample size, the SD of each combination of two miRNAs is calculated and averaged. This process is repeated 20 times, and the error bars show the standard deviation of the repeats. The weights were calculated based on Raw CT values. (D) The number of reference genes effect on stability. For each number of reference genes, the SD of different combinations of miRNAs were calculated. SD: Standard Deviation, Normalized: weights were optimized on the normalized data, raw CT: weights were optimized on the Raw CT values, geom: usual geometric mean

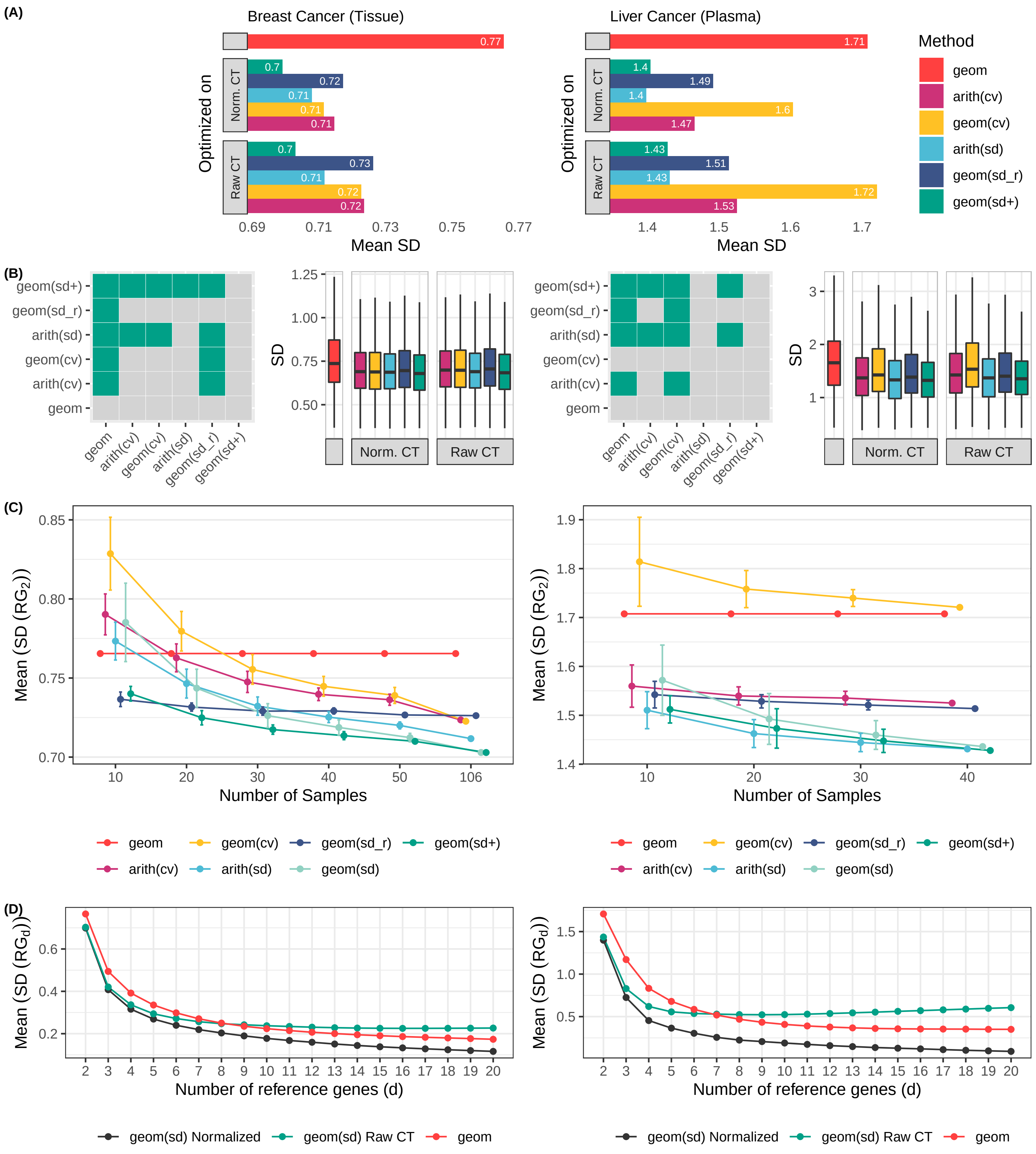


Figure S.6: Stability (SD) comparison of different weight methods. Figures on the left and right side are for the breast cancer and liver cancer respectively. (A) Mean stability of all combinations of two miRNAs in different weight methods (the lower is better). (B) Box plots for the stability of all combinations of two miRNAs in different weight methods and the tile figures show paired Wilcoxon test between the stability of different weight methods on Raw CT values. Colored tiles indicate that the row weight method had significantly ($p<0.01$) lower stability than the column one. (C) Sample size analysis: for each sample size, the SD of each combination of two miRNAs is calculated and averaged. This process is repeated 20 times, and the error bars show the standard deviation of the repeats. The weights were calculated based on Raw CT values. (D) The number of reference genes effect on stability. For each number of reference genes, the SD of different combinations of miRNAs were calculated. SD: Standard Deviation, Normalized: weights were optimized on the normalized data, raw CT: weights were optimized on the Raw CT values, geom: usual geometric mean

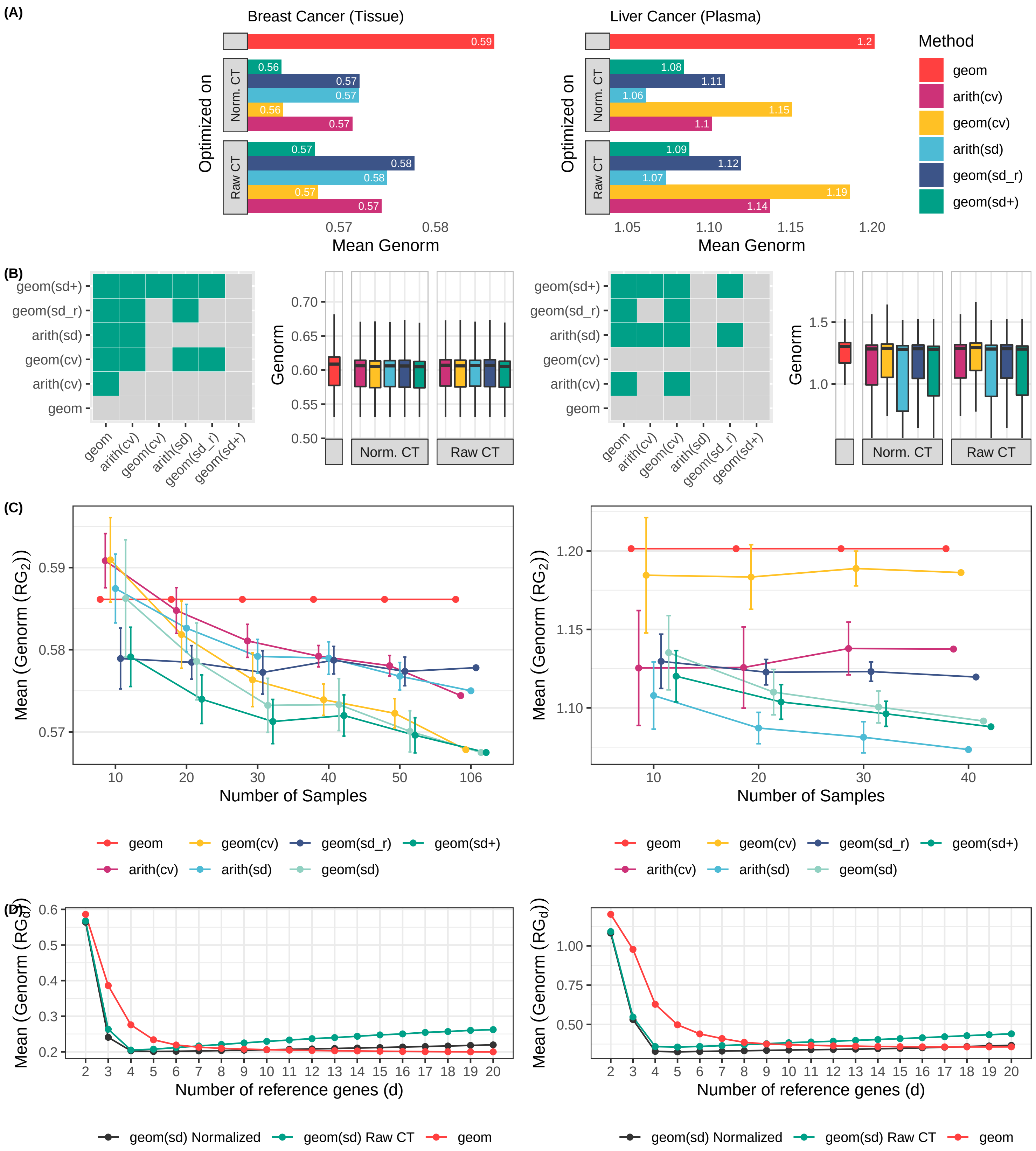


Figure S.7: Stability (Genorm score) comparison of different weight methods (lower is better). Figures on the left and right side are for the breast cancer and liver cancer respectively. (A) Mean stability of all combinations of two miRNAs in different weight methods (the lower is better). (B) Box plots for the stability of all combinations of two miRNAs in different weight methods and the tile figures show paired Wilcoxon test between the stability of different weight methods on Raw CT values. Colored tiles indicate that the row weight method had significantly ($p<0.01$) lower stability than the column one. (C) Sample size analysis: for each sample size, the SD of each combination of two miRNAs is calculated and averaged. This process is repeated 20 times, and the error bars show the standard deviation of the repeats. The weights were calculated based on Raw CT values. (D) The number of reference genes effect on stability. For each number of reference genes, the SD of different combinations of miRNAs were calculated. SD: Standard Deviation, Normalized: weights were optimized on the normalized data, raw CT: weights were optimized on the Raw CT values, geom: usual geometric mean

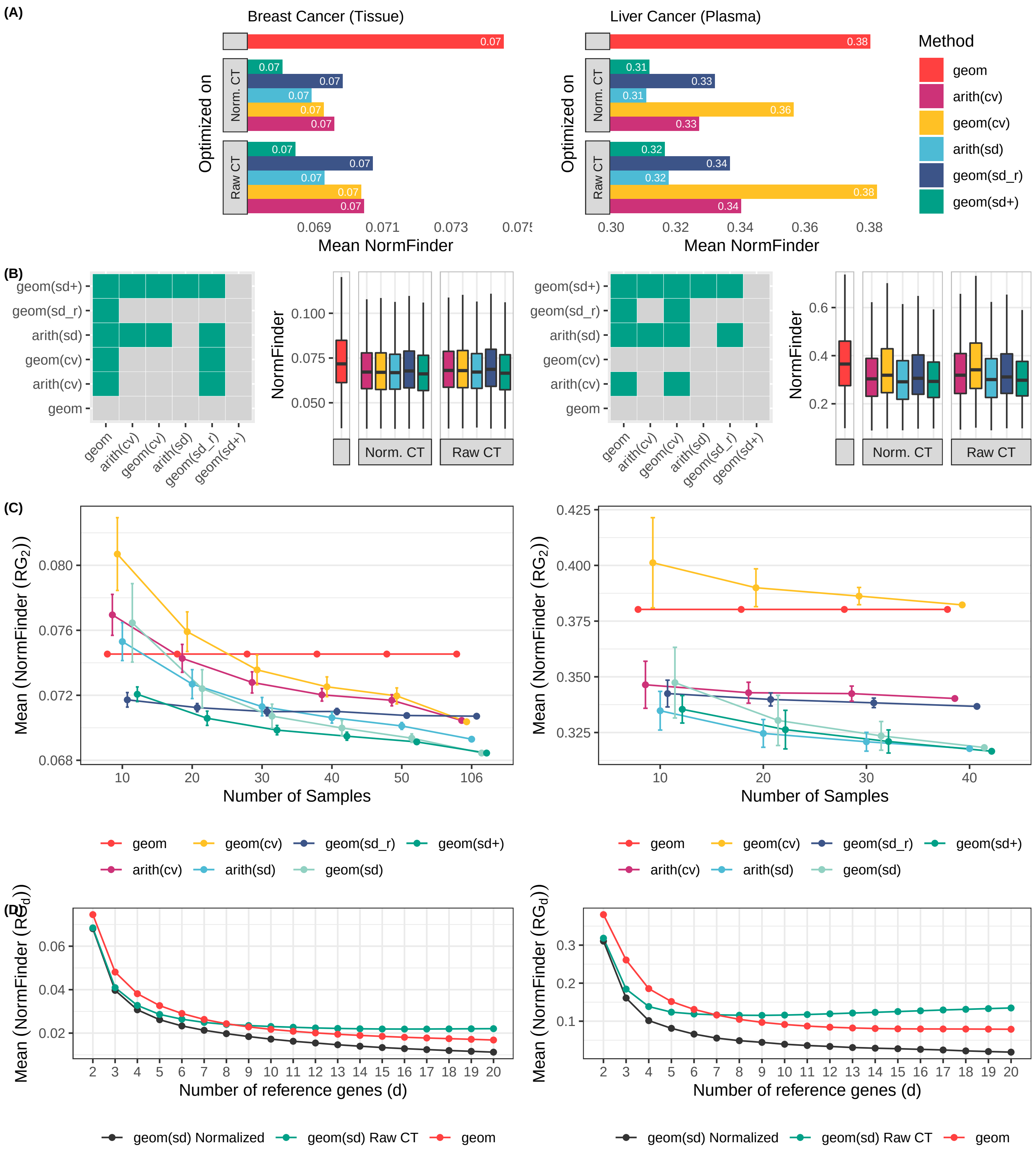


Figure S.8: Stability (NormFinder score) comparison of different weight methods (lower is better). Figures on the left and right side are for the breast cancer and liver cancer respectively. (A) Mean stability of all combinations of two miRNAs in different weight methods (the lower is better). (B) Box plots for the stability (NormFinder score) of all combinations of two miRNAs in different weight methods and the tile figures show paired Wilcoxon test between the stability of different weight methods on Raw CT values. Colored tiles indicate that the row weight method had significantly ($p<0.01$) lower stability than the column one. (C) Sample size analysis: for each sample size, the SD of each combination of two miRNAs is calculated and averaged. This process is repeated 20 times, and the error bars show the standard deviation of the repeats. The weights were calculated based on Raw CT values. (D) The number of reference genes effect on stability. For each number of reference genes, the SD of different combinations of miRNAs were calculated. SD: Standard Deviation, Normalized: weights were optimized on the normalized data, raw CT: weights were optimized on the Raw CT values, geom: usual geometric mean

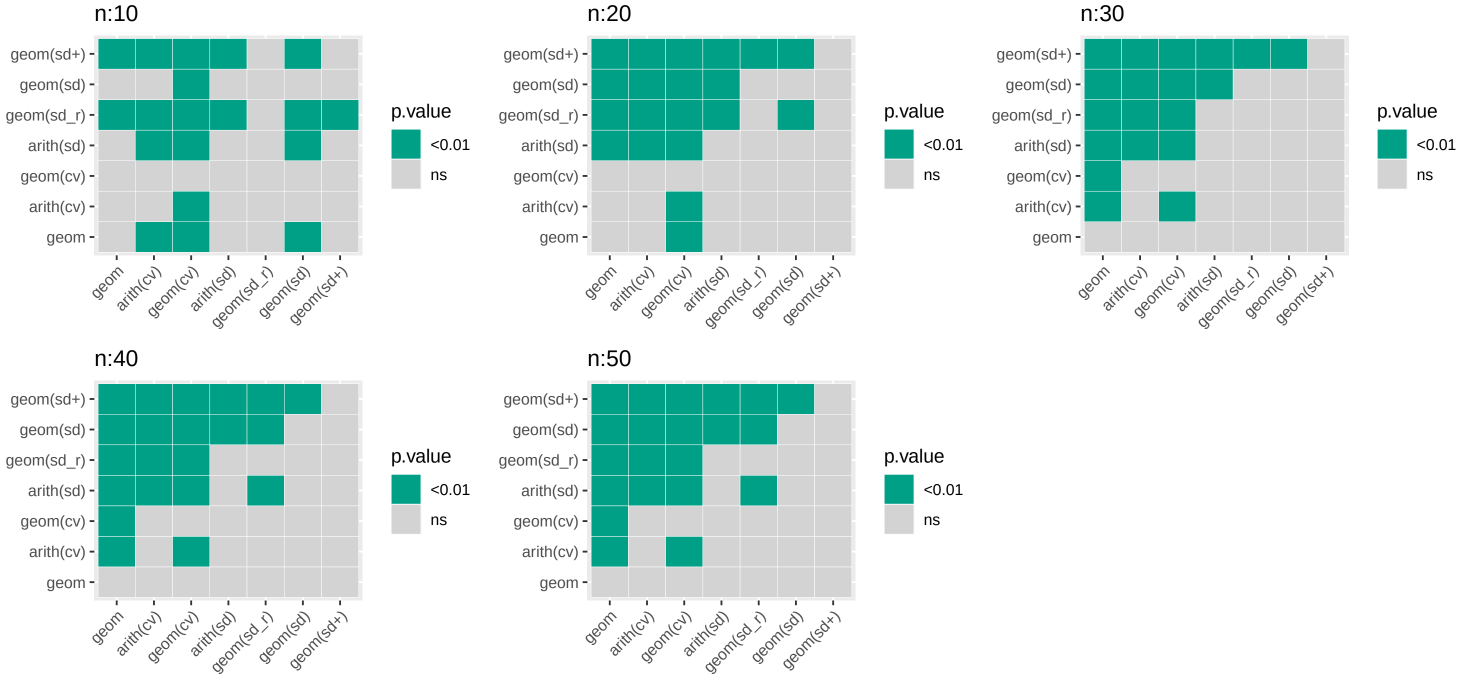


Figure S.9: Statistical stability comparison of weighting methods in different sample sizes (n) on GSE78870 breast cancer dataset. Each tile plot shows paired Wilcoxon test between the stability of different weighting methods on raw CT values. Colored tiles indicate that the row weighting method had significantly ($p<0.01$) lower stability than the column one. each sub-sampling was repeated 20 times, with the average of the results considered. This figure is complements the Fig2C of the paper.

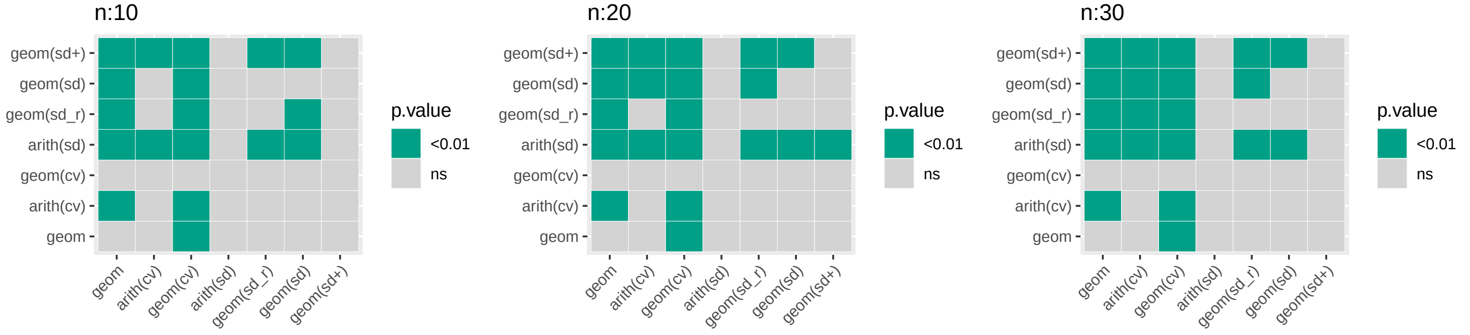


Figure S.10: Statistical stability comparison of weighting methods in different sample sizes (n) on GSE50013 liver cancer dataset. Each tile plot shows paired Wilcoxon test between the stability of different weighting methods on raw CT values. Colored tiles indicate that the row weighting method had significantly ($p<0.01$) lower stability than the column one. each sub-sampling was repeated 20 times, with the average of the results considered. This figure is complements the Fig2C of the paper.
